# A Unique Promoter Architecture Defines Menin-KMT2A Complex Selectivity

**DOI:** 10.64898/2026.10.30.763078

**Authors:** Pablo R. Freire, Brandon Regalado, Reina Takeda, Elizabeth T. Ener, George Wan, Jevon A. Cutler, Tavor Z. Baharav, Jared Brown, Vassilena Sharlandjieva, Thomas A. Milne, Rafael A. Irizarry, Scott A. Armstrong

## Abstract

Selective occupancy of chromatin-regulatory complexes is central to developmental gene regulation, yet the sequence determinants of this selectivity remain poorly understood. Using interpretable machine learning, we identify a composite promoter architecture comprising CpG islands flanked by several-kilobase A/T-rich domains (AT-flanks) that denotes high occupancy of the Menin-KMT2A histone-modifying complex at developmental and leukemia-associated genes. Genetic deletion and chromatin-contact analyses show that AT-flanks support KMT2A retention at linked CpG islands. We identify LEDGF as an A/T-rich DNA-associated protein required for Menin engagement at AT-flanks and sustained KMT2A promoter occupancy. AT-flank extent is associated with differential KMT2A loss following pharmacologic disruption of the Menin-KMT2A interaction, providing a genomic basis for heterogeneous target sensitivity to recently approved Menin inhibitors in acute leukemia. Beyond Menin-KMT2A, the same architecture distinguishes promoters with high canonical PRC1 occupancy across diverse cell types, revealing a shared sequence organization associated with distinct chromatin-regulatory states.

## Introduction

Chromatin-based regulation of gene expression is essential for development and cellular identity. The discovery of Polycomb group (PcG) and trithorax group (TrxG) genes through homeotic transformations in *Drosophila* established a foundational model in which opposing chromatin regulators maintain repressed or active states of homeotic (Hox) genes after their initial expression patterns have been established^1^. Mammals encode two co-orthologs of *Drosophila* trithorax, KMT2A and KMT2B (lysine methyltransferases 2A and 2B, also known as MLL1 and MLL2), which function within a COMPASS (Complex of Proteins Associated with Set1)-like assembly^2^ and contribute to the deposition of histone H3 lysine 4 trimethylation (H3K4me3) at a subset of active gene promoters^3,4^, including developmental loci such as Hox genes. COMPASS-like complexes share a conserved catalytic core complex that associates with the C-terminal SET domain and supports H3K4 methylation, while individual KMT2 family members engage distinct additional interactors that correlate with differences in genomic distribution and biological function^2,3^. In particular, KMT2A’s biological activity extends beyond H3K4 methylation, and mice lacking its SET domain survive into adulthood, whereas germline deletion of *Kmt2a* is embryonic lethal^5,6^. Importantly, *KMT2A* is the target of recurrent chromosomal translocations in acute leukemia that fuse its N-terminal region to diverse partner proteins primarily involved in transcriptional elongation, resulting in fusion proteins that lack the C-terminal SET domain^7–9^.

Genome-wide studies identified human KMT2A at the promoter-proximal regions of thousands of actively transcribed genes^10^. KMT2A associates with distinct regulatory factors, including Menin and host cell factor 1 (HCF-1)^11^, which can mark different classes of KMT2A-bound promoters. Whereas HCF-1 is broadly associated with housekeeping and cell-cycle-regulated genes^12^, Menin occupancy is highly concentrated at a subset of KMT2A-associated promoters, including developmental and cell-cycle regulators such as *HOX* genes, *MEIS1*, and cyclin-dependent kinase inhibitors^13–15^. At these loci, Menin and KMT2A associate with the PWWP-domain chromatin reader LEDGF (Lens Epithelium-derived Growth Factor)^16^.

In hematopoiesis, Menin is essential for the repopulating capacity of stem cells, while being dispensable for steady-state hematopoiesis^17^. Genetically distinct acute leukemias driven by *KMT2A* rearrangements (*KMT2A-r*), *NPM1c* mutations, or *NUP98* rearrangements converge on a Menin-KMT2A-dependent transcriptional program characterized by sustained expression of developmental regulators, most prominently *HOX* genes and *MEIS1*, which promotes aberrant progenitor self-renewal and impairs differentiation^18–23^. Detailed studies of the Menin-KMT2A complex have shown that Menin interacts directly with KMT2A through a structurally defined binding domain on Menin that can be targeted by multiple potent small molecule inhibitors^24,25^. Disruption of this protein-protein interaction leads to reversal of gene expression, cellular differentiation and ultimately leukemia cell death^14,15,26^. These small molecule Menin inhibitors (MENi) have shown remarkable efficacy against acute myeloid leukemia (AML) in clinical trials^27–29^, culminating in their FDA approval for the treatment of relapsed or refractory *KMT2A*-r and *NPM1c*-mutated leukemia. Despite extensive structural and therapeutic characterization of the Menin-KMT2A-LEDGF interactions, the mechanistic role of Menin and LEDGF within the KMT2A complex remains poorly defined. Furthermore, Menin inhibition affects only a subset of Menin-KMT2A target genes, indicating unrecognized heterogeneity in promoter dependence on this interaction^14,15^.

The developmentally important promoters selectively bound by Menin-KMT2A, including *HOX* genes and *MEIS1*, are often prominent targets of mammalian Polycomb complexes. In *Drosophila*, PcG and TrxG regulation can converge on sequence-encoded PRE/TRE elements^1^, but the corresponding DNA-encoded determinants of mammalian KMT2A and Polycomb occupancy remain incompletely defined^30^. CpG islands provide an important recruitment platform for both KMT2A^31,32^ and Polycomb complexes^33–36^, while discrete DNA elements capable of tethering canonical Polycomb Repressive Complex 1 (cPRC1) have also been identified^37^. Nevertheless, CpG-island recognition alone cannot explain why Menin-KMT2A or Polycomb complexes become concentrated at restricted subsets of predominantly developmental promoters.

Here, we use an interpretable machine learning approach to uncover a promoter architecture in which large A/T-rich domains flanking CpG islands specify Menin-KMT2A occupancy. These A/T-rich flanks (AT-flanks) engage in broad chromatin interactions and function as genomic elements that are distinct from canonical enhancers, with AT-flank abundance correlating with promoter sensitivity to Menin inhibition. We further identify LEDGF as a direct AT-flank-binding factor that contributes to Menin-KMT2A retention at these loci. Extending this observation, the same promoter architecture is associated with high cPRC1 occupancy across multiple cell lines, with cPRC1 members preferentially occupying AT-flanks. Our results establish AT-flanks as promoter elements that support locus-selective Menin-KMT2A occupancy and as a broader genomic feature associated with canonical PRC1 binding.

## Results

### A/T-rich TSS-flanks are associated with Menin-KMT2A occupancy

In mammals, KMT2A binds to a large number of promoter-associated CpG islands in combination with different complex members such as Menin and HCF-1^11^, which occupy mutually exclusive gene sets (Figure S1A,B). While HCF-1 is commonly present at promoters of housekeeping genes^12^, Menin is disproportionately associated with a small number of promoters in both cancer and non-transformed human primary cells (Figure S1C). At these promoters, KMT2A and Menin have a broad binding pattern, extending several kilobases upstream and downstream of the transcriptional start site (TSS) (Figure S1D). Importantly, analysis of known and de novo DNA motifs using traditional motif search algorithms failed to identify unique sequences associated with high Menin-KMT2A occupancy at promoters (Figure S2A). These observations led us to hypothesize that a more complex DNA sequence pattern, beyond the sole presence/absence of short transcription factor binding sites, may be associated with the asymmetrically high occupancy of the Menin-KMT2A complex at a select set of loci.

To uncover this pattern, we implemented a convolutional neural network (CNN) binary classifier to predict promoters with disproportionately high Menin-KMT2A occupancy using a large −3 to +20 kb sequence window around the TSS as the sole input (Figure 1A). The choice of the window size was informed by the broad occupancy of Menin and KMT2A at the TSS. CNNs use filters (“kernels”) that are capable of detecting complex DNA motifs of arbitrary lengths and multiplicity, while accounting for potential interactions between them^38^. Unlike many state-of-the-art DNA machine learning algorithms^39,40^, we opted for a simpler CNN architecture with only two convolutional layers to facilitate model interpretation and the extraction of biological insight. Training and validation sets consisted of promoter sequences with high Menin and KMT2A occupancy (Menin-KMT2A^high^) or low Menin occupancy (Menin^low^) in the AML cell lines MV4-11 (*KMT2A::AFF1* fusion) and OCI-AML3 (*NPM1c*-mutant, wild-type *KMT2A*). Model performance was evaluated on an independent, non-overlapping test set derived from HeLa cells, achieving a ROC AUC of 0.89 (Figure 1B). The use of a smaller −3 to +3 kb window resulted in reduced performance (Figure S2B), suggesting that unique sequences several kilobases upstream and downstream of the TSS are correlated with high Menin-KMT2A occupancy.

**Figure 1.**
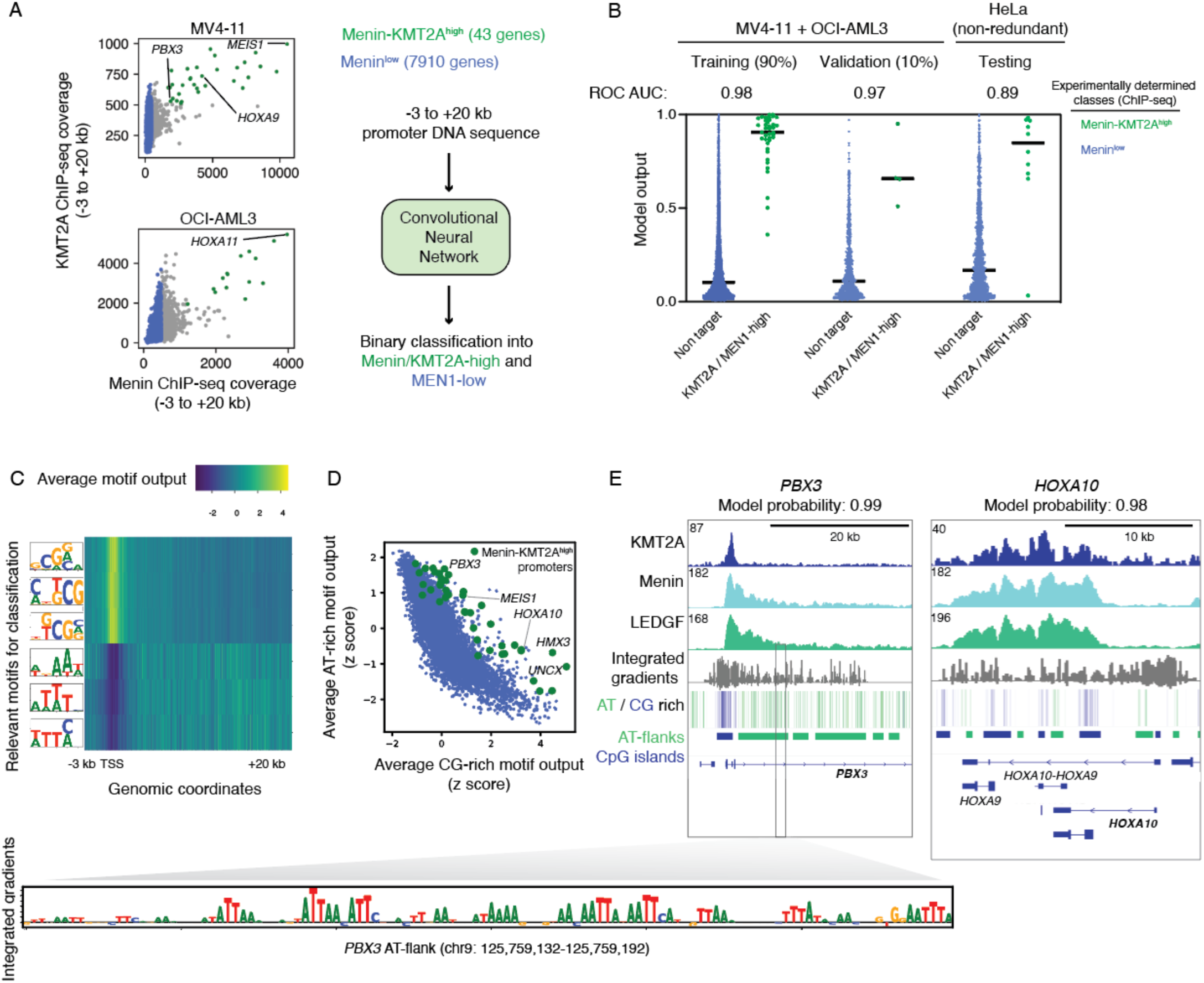
Menin-KMT2A occupancy is associated with A/T-rich flanks. **a,** Schematic model of the CNN classifier and training/validation class definition based on KMT2A and Menin ChIP-seq data from MV4-11 and OCI-AML3 cells. **b**, CNN model output. **c,** First-layer kernel outputs on the -3 to +20 kb region averaged across Menin-KMT2Ahigh promoters. **d,** Total kernel outputs from C/G- and A/T-rich motifs (averaged across -3 to +20 kb) in Menin-KMT2A^high^ and Menin^low^ promoters in the training and validation sets. **e,** Integrated gradient analysis of the contribution of input nucleotides to the overall classification, overlaid with ChIP-seq data for KMT2A, Menin, and LEDGF in MV4-11 cells. ROC, receiver operating characteristic; AUC, area under the curve.

To identify the specific DNA motifs associated with Menin-KMT2A occupancy, we measured the contribution (integrated gradient) of each kernel to the final output. We found that a simple combination of 6 A/T- and C/G-rich motifs was sufficient for accurate classification of Menin-KMT2A-occupied TSS regions (Figure 1C). In most human promoters, the frequencies of these C/G- and A/T-rich motifs are inversely correlated. However, at Menin-KMT2A^high^ targets, both A/T- and C/G-rich motifs are enriched within the same promoter (Figure 1D), revealing a unique architecture consisting of CpG islands flanked by A/T-rich regions (AT-flanks) extending several kilobases away from the TSS. We next assessed the contribution of each individual nucleotide to the final output, revealing that broad regions encompassing both CpG islands and AT-flanks contributed to the correct classification by the CNN model (Figure 1E).

To establish the simplest and smallest DNA motifs associated with Menin-KMT2A occupancy, we next implemented a logistic regression classifier using DNA k-mer counts of different lengths and in varying input windows. The simple count of di-nucleotides from -4 to +12 kb was sufficient for the effective classification of Menin-KMT2A^high^ targets (5-fold cross-validation ROC AUC > 0.95) (Figure S3). Increasing the k-mer length and input window size only marginally improved classification. Accordingly, the CG, AA and AT di-nucleotides had the highest regression coefficients (Table S1). Together, these findings reveal a sequence architecture distinct from conventional motif-centered transcription-factor recognition, in which information associated with Menin-KMT2A occupancy is distributed across several-kilobase promoter domains.

Notably, the ratio between AT-flanks and CpG islands varied among Menin-KMT2A targets. Some genes, such as *PBX3* and *MEIS1*, have large AT-flanks and smaller CpG islands, whereas others, including *HOXA* factors and *BAHCC1*, have extensive CpG islands interspersed with smaller AT-flanks (Figure S4). In Menin-KMT2A^high^ targets with large AT-flanks, Menin and its binding partner LEDGF occupied both CpG islands and AT-flanks, whereas KMT2A was restricted to the CpG islands. In contrast, targets with small AT-flanks had greater co-localization of KMT2A, Menin and LEDGF throughout their promoters. Despite these differences, the co-occurrence of CpG islands and AT-flanks was disproportionately high in all Menin-KMT2A^high^ targets when compared to Menin^low^ promoters (Figure 1D).

Overall, Menin-KMT2A^high^ AT-flanks were more conserved than canonical enhancers but lacked the typical enhancer features such as open chromatin and high occupancy of BRD4, CDK8 and H3K27ac^41,42^, as illustrated by comparing the Menin-KMT2A^high^ locus *PBX3* to the Menin^low^ locus *SPI1* (Figure S5A and B). When compared to the candidate cis-regulatory elements defined by the ENCODE project^43^, AT-flanks at Menin-KMT2A^high^ promoters were more conserved than distal and proximal enhancer elements but less conserved than CpG islands and promoter regions in general (Figure S5C). Remarkably, AT-flanks present in Menin^low^ promoters were less conserved than all other tested elements, indicating a potential conserved functional role for AT-flanks at Menin-KMT2A^high^ targets.

### KMT2A Loss Following Menin Inhibition Correlates with AT-Flank Extent

Previous studies have shown that MENi have variable effects on the expression of Menin-KMT2A target genes, with some, such as *MEIS1,* being consistently suppressed in cell lines and patient samples, whereas others, such as *HOXA* genes, have variable responses^14,15,26,44^. Confirming previous results, we found that MENi treatment in AML cell lines carrying either *KMT2A-r* (MV4-11 and MOLM13) or wild-type *KMT2A* (OCI-AML3) caused the eviction of Menin from chromatin, yet KMT2A remained partially retained at the CpG islands of select genes, including *HOXA* factors and *BAHCC1* (Figure 2A). Globally, MENi treatment resulted in a variable reduction of KMT2A occupancy at Menin-KMT2A^high^ targets (Figure 2B). Notably, the extent of KMT2A loss correlated with the size of AT-flanks surrounding the TSS (Figure 2C). Genes with large Menin-bound AT-flanks, such as *MEIS1* and *PBX3*, exhibited significant KMT2A loss upon MENi treatment, whereas promoters with small AT-flanks, including *HOXA10* and *UNCX*, showed more modest reductions, or even an increase in KMT2A binding in the case of *BAHCC1.* Loss of KMT2A at promoters was significantly correlated with reduction in gene expression, although, in some cases, genes retained similar expression despite the loss of KMT2A (Figure S6).

**Figure 2.**
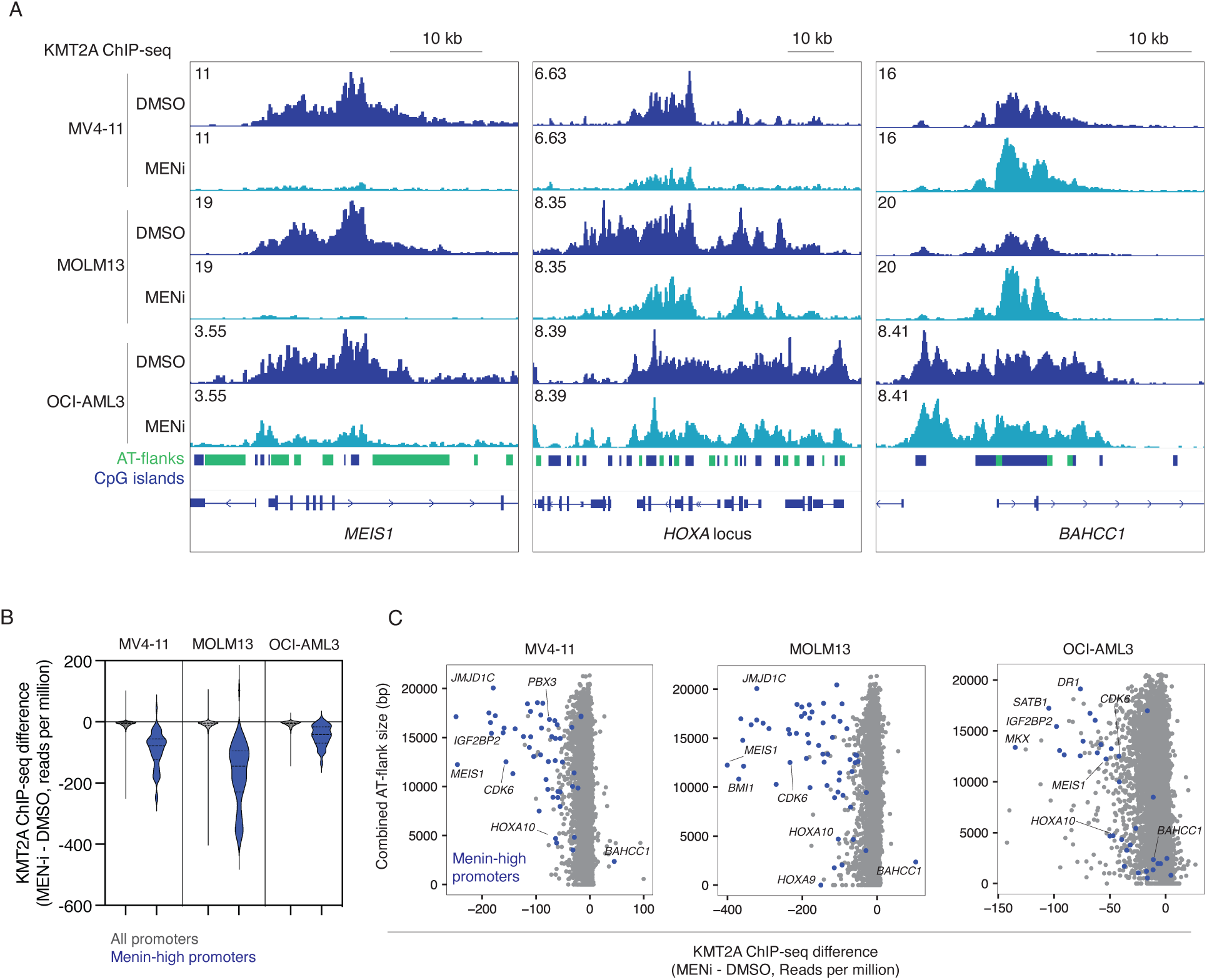
AT-flank sizes are associated with the response to Menin inhibition. **a,** KMT2A ChIP-seq tracks at Menin-KMT2A^high^ targets in MV4-11, MOLM13 and OCI-AML3 cells after treatment with Menin inhibitor (VTP-50469, 500 nM, 3 days). **b,** Changes in KMT2A occupancy after Menin inhibition. Dashed line: median, dotted line: quartiles. **c,** Scatter plot showing the relationship between combined AT-flank sizes (-3 to +20 kb window) and KMT2A occupancy changes upon Menin inhibition.

Together, our results indicate that Menin-KMT2A target genes can be broadly divided into two classes: one class characterized by large AT-flanks that is more likely to lose KMT2A occupancy upon Menin inhibition, and a second class with smaller AT-flanks that preferentially retains KMT2A following MENi treatment. In the latter group, KMT2A recruitment may be mediated predominantly through CpG islands, potentially via direct DNA binding by its CXXC domain in a Menin-independent manner^32,45^. Consistent with this model, at Menin-inhibitor-sensitive promoters, Menin and its binding partner LEDGF broadly occupy large AT-flank regions whereas KMT2A is restricted to the CpG island, while at less sensitive promoters, Menin, LEDGF, and KMT2A exhibit greater co-occurrence across the promoter region (Figure S4).

### AT-Flanks Sustain Menin-KMT2A Occupancy and Active Promoter Chromatin

To assess the functional role of AT-flanks in regulating chromatin state and expression of Menin-KMT2A^high^ genes, we generated homozygous CRISPR/Cas9 deletions in MV4-11 cells targeting a downstream 22 kb AT-flank at the *PBX3* locus and an upstream 2.4 kb AT-flank at the *CDK6* locus. AT-flank deletion led to the eviction of KMT2A from the nearby CpG island despite the > 800 bp separation between the deleted regions and the CpG island (Figure 3A). Furthermore, the presence of Menin, LEDGF and ENL (an elongation factor associated with KMT2A^46,47^) at promoters was significantly reduced. AT-flank deletions also led to a decrease in RNA Pol II occupancy and reduced levels of the activating histone modifications H3K9ac and H3K27ac, whereas the levels of H3K4me3 remained mostly unchanged. AT-flank deletions caused a reduction in gene expression of both *PBX3* and *CDK6* (Figure 3B). To examine the impact of AT-flank size on the activity of the Menin-KMT2A complex, we generated a smaller 8.6 kb deletion of the *PBX3* AT-flank, which caused a less pronounced reduction in gene expression compared to the larger 22 kb deletion (Figure 3C), suggesting that AT-flanks contribute to Menin-KMT2A activity in a length-dependent manner. Additionally, BRD4 ChIP-seq and ATAC-seq revealed the presence of a putative 2 kb enhancer at the center of the 22 kb *PBX3* AT-flank. However, its deletion did not alter gene expression or Menin occupancy (Figure 3C). Finally, to rule out general transcriptional effects of promoter flanking regions, we deleted the flanking regions of two Menin^low^ target genes, *SMARCA4* and *CD47*, and observed no significant changes in their expression (Figure S7).

**Figure 3.**
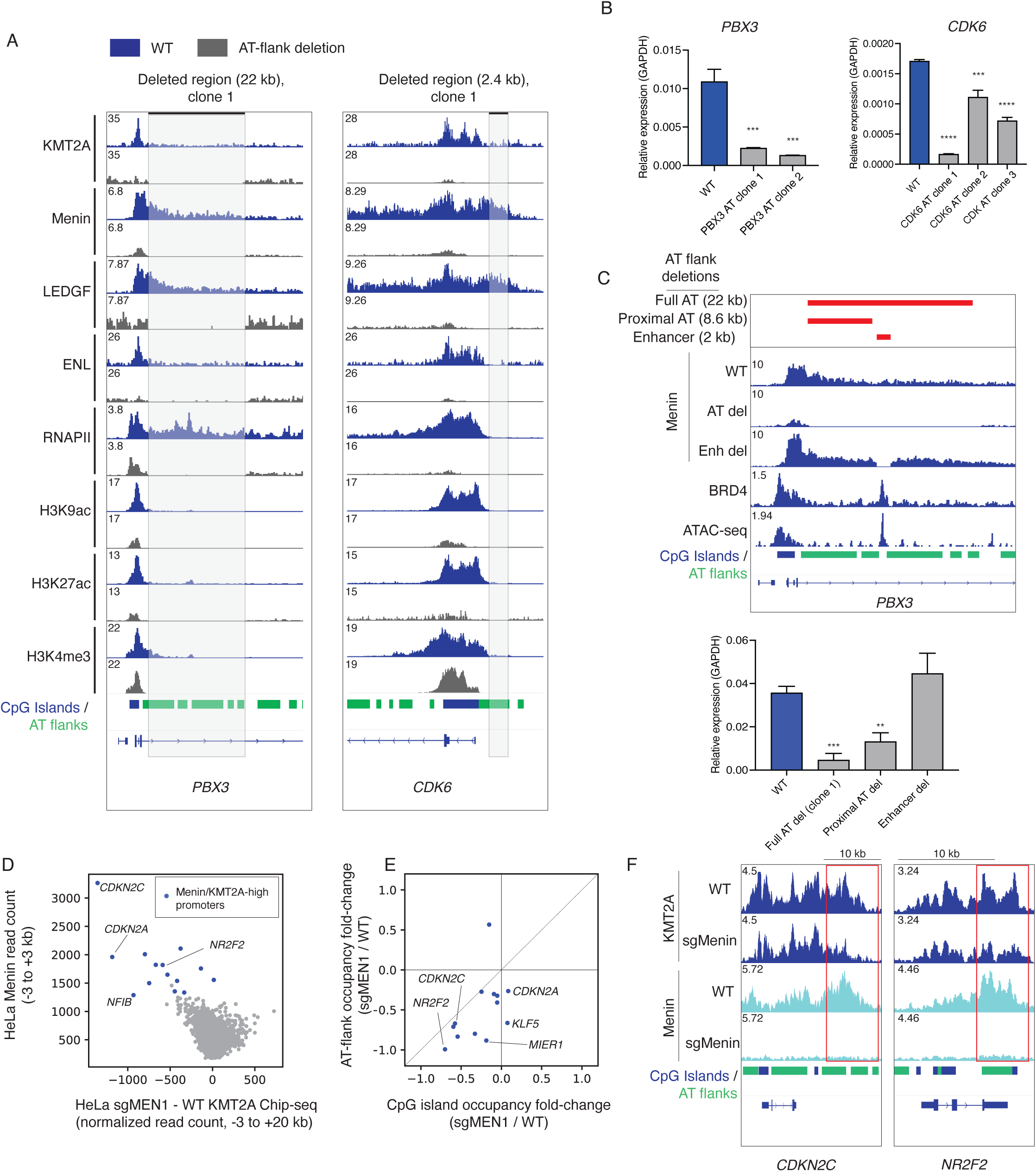
AT-flanks sustain KMT2A occupancy and gene expression. **a,** ChIP-seq analysis of KMT2A, Menin, LEDGF, ENL, RNA Pol II, H3K9ac, H3K27ac, and H3K4me3 after CRISPR/Cas9-mediated AT-flank deletion at *PBX3* and *CDK6* in MV4-11 cells. *b,* RT-qPCR analysis of *PBX3* and *CDK6* expression after AT-flank deletion. **c,** Effects of partial *PBX3* AT-flank deletions on Menin occupancy (ChIP-seq) and gene expression (RT-qPCR). **d**, Scatter plot comparing Menin occupancy and KMT2A loss after Menin depletion in HeLa cells. **e,** Comparison of KMT2A loss at CpG islands and AT-flanks upon Menin depletion in HeLa cells. The diagonal line represents the identity line (x = y). **f,** Representative changes in KMT2A and Menin ChIP-seq occupancy in HeLa cells upon Menin depletion. Error bars indicate s.d.; P-values were calculated using a two-tailed t-test, n=3, **P < 0.01, *** P<0.001, **** P<0.0001 (b and c). KMT2A ChIP-seq coverage data were normalized with median ratio normalization (d and e).

As an orthogonal approach, we examined Menin’s role in AT-flank-mediated KMT2A association using HeLa cells, which express wild-type KMT2A and are not dependent on the Menin-KMT2A complex for proliferation, enabling the long-term deletion of Menin. HeLa cells also exhibit a distinctive ChIP-seq profile in which KMT2A occupies both CpG islands and AT-flanks, whereas Menin is more strongly enriched at AT-flanks. This pattern allows the direct assessment of the involvement of AT-flanks in KMT2A occupancy. Menin depletion caused the displacement of KMT2A at Menin-KMT2A^high^ promoters (Figure 3D), and this effect was particularly strong at AT-flanks (Figure 3E and F). Together, these results support a functional role for AT-flanks in sustaining Menin-KMT2A and RNA polymerase II occupancy, as well as active promoter chromatin, at Menin-KMT2A^high^ target genes.

### AT-flanks Form Broad Contacts with Linked CpG Islands

The observation that AT-flank deletions disrupt KMT2A recruitment to CpG islands > 800 bp away (Figure 3A) prompted us to investigate the three-dimensional genomic architecture at Menin-KMT2A^high^ promoters. To this end, we first performed Micro Capture-C (MCC)^48^ on four Menin-KMT2A^high^ target genes with large AT-flanks (*CDK6*, *MEIS1*, *FOXP1* and *IGF2BP2*), in which KMT2A recruitment is sensitive to MENi treatment. MCC enables high-resolution mapping of chromatin contacts between selected genomic viewpoints (“anchors”) and surrounding regions, allowing the detection of fine-scale promoter-associated interactions. Using both CpG islands and AT-flanks as capture anchors, we detected extensive long-range interactions between these elements that spanned several kilobases on either side of the transcription start site (Figure 4A,B). Unlike canonical enhancer-promoter loops^49^, these interactions lacked a discrete interaction peak and instead appeared as diffuse contact enrichments across the AT-flank regions. Notably, treatment with MENi disrupted AT-flank-CpG island contacts, highlighting the role of the Menin-KMT2A interaction in maintaining this promoter 3D configuration (Figure 4A). Furthermore, treatment of cells with triptolide, an RNA Pol II initiation inhibitor^50^ (Figure S8A), did not affect the interactions between CpG islands and AT-flanks, indicating that ongoing transcription is not required for the maintenance of AT-flank chromatin contacts. Our MCC analysis also detected concomitant interactions between CpG islands and canonical enhancers at the *CDK6* and *FOXP1* loci, as evidenced by ATAC-seq and BRD4 ChIP-seq (Figure 4B). However, treatment with MENi selectively disrupted CpG island-AT-flank interactions, while leaving CpG island-enhancer interactions intact (Figure S8B).

**Figure 4.**
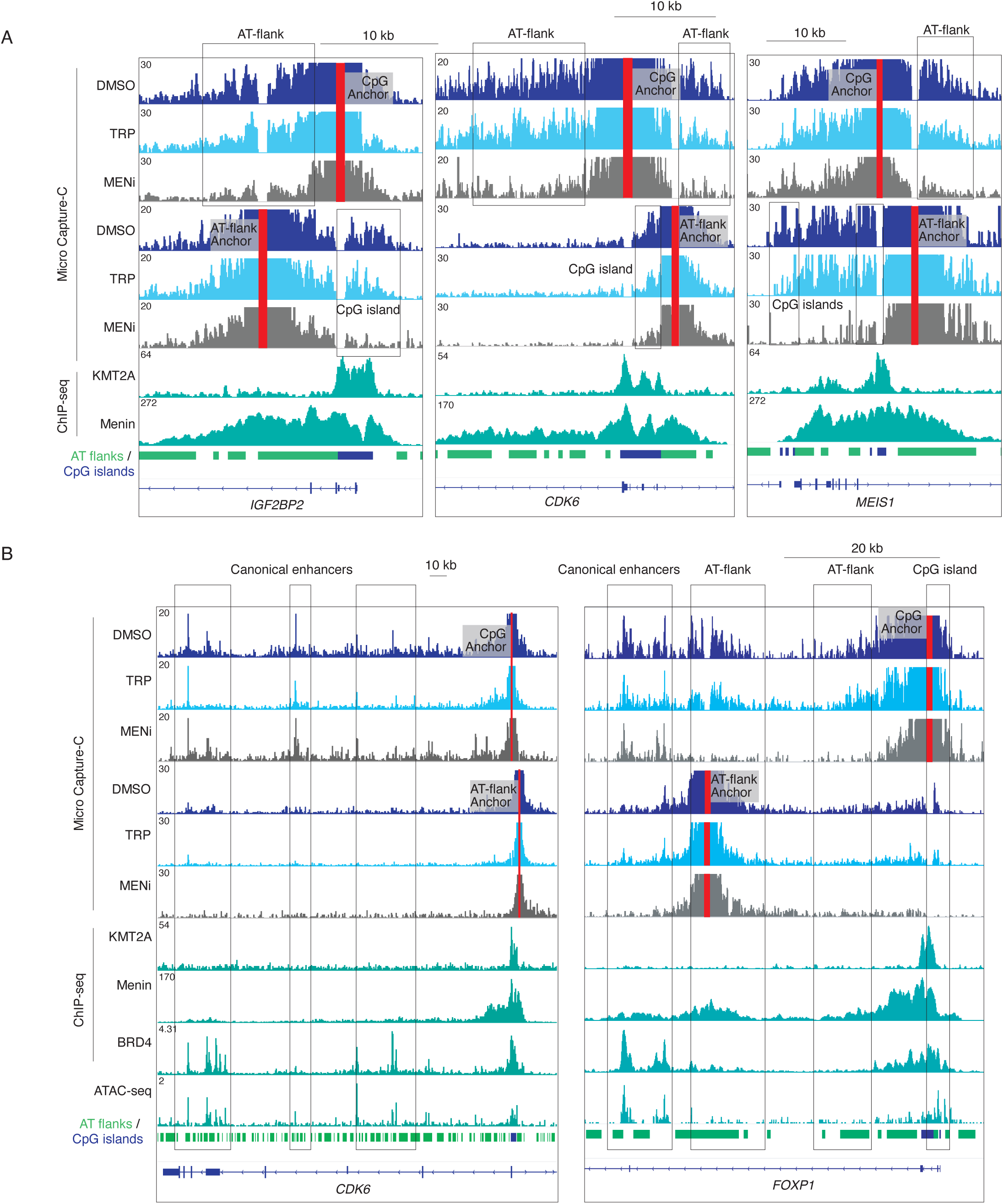
AT-flanks form long-range interactions with CpG islands. Micro Capture-C (MCC) analysis in MV4-11 cells upon Menin inhibition (VTP-50469, 500 nM, 3 days) and RNA Pol II initiation inhibition (triptolide, 500 nM, 1 h). ChIP-seq tracks for KMT2A and Menin are shown. Red boxes indicate captured regions (CpG island or AT-flank anchors). **a,** MCC analysis on the *IGF2BP2*, *CDK6* and *MEIS1* loci. **b,** Micro Capture-C analysis on *CDK6* and *FOXP1* loci along with ATAC-seq and BRD4 ChIP-seq data to mark canonical enhancers. MENi, Menin inhibitor; TRP, triptolide.

We next performed MCC on Menin-KMT2A^high^ targets with small AT-flanks (*HOXA* locus and *BAHCC1*), in which KMT2A recruitment is less sensitive to MENi treatment. At the *HOXA* locus, MENi treatment induced a small reduction in the interactions adjacent to the probed locus, but distal interactions were not observed (Figure S8C, D). In contrast, at *BAHCC1*, MENi treatment had no effect on 3D interactions between the CpG islands and adjacent regions. Collectively, these results show that AT-flanks engage in broad long-range contacts with linked CpG islands that depend on the Menin-KMT2A interaction. The broad topology of these contacts, together with their differential sensitivity to MENi relative to most distal enhancer-promoter interactions, supports the conclusion that AT-flank-CpG island contacts represent a distinct mode of promoter-associated chromatin organization.

### LEDGF preferentially binds to A/T-rich DNA in vitro

The presence of biochemically validated DNA- and histone-binding domains in LEDGF prompted us to investigate whether it engages AT-flanks through direct DNA contact. In particular, LEDGF contains AT-hook-like motifs, which have been shown to bind the minor groove of A/T-rich DNA sequences in other proteins^51^. Previous studies have shown that the AT-hook domains of LEDGF mediate DNA binding in vitro and exhibit an intrinsic affinity for DNA even in the absence of other domains^52,53^. Additionally, an N-terminal PWWP domain recognizes H3K36me2/3 and contacts nucleosomal DNA through adjacent regions^54,55^, although it lacks a strong affinity for naked DNA in vitro^52,53^. More recent NMR and biochemical analyses further demonstrated that a flexible region encompassing the AT-hook-containing segment directly contacts DNA, supporting a multivalent binding mode in which DNA engagement is distributed across several LEDGF regions^56^. Despite this extensive evidence for direct DNA engagement, no defined DNA sequence preference has been described for LEDGF, and whether this multivalent architecture confers selectivity for particular genomic sequence contexts has remained unclear.

To assess the affinity of LEDGF for DNA in vitro, we used full-length LEDGF and domain deletion mutants expressed in *E. coli* (Figure 5A). To test whether LEDGF exhibits sequence bias in DNA binding in vitro, we first performed a fluorescence polarization (FP) assay using recombinant full-length LEDGF (Figure 5B), which showed that LEDGF binds to the AT-flank region of *IGF2BP2* in vitro, while no binding could be detected to probes derived from the *IGF2BP2* CpG island or a *GAPDH* intron that is neither A/T nor C/G rich. Notably, we did not detect binding of recombinant full-length Menin to any of the tested DNA probes. We next measured the binding of LEDGF to artificial probes designed to mimic the CpG islands (CG-probes, high CpG count) and AT-flank regions (AT-probes, high AAA/TTT counts) using electrophoretic mobility shift assay (EMSA), which was more sensitive than FP in detecting LEDGF-DNA interactions. In the absence of unlabeled competitors, LEDGF was capable of binding both AT- and CG-probes (Figure 5C). However, the addition of a 20-fold excess of unlabeled AT-probe abolished LEDGF binding to labeled CG-probe, while the presence of a 20-fold excess of unlabeled CG-probe did not affect LEDGF binding to biotinylated AT-probe, suggesting that LEDGF preferentially binds A/T-rich sequences. The addition of unlabeled competitors with the same sequence markedly reduced LEDGF binding to both AT- and CG-probes, confirming that both unlabeled probes effectively competed for LEDGF binding under the assay conditions. Furthermore, a 20-fold excess of an *IGF2BP2* AT-flank unlabeled probe significantly reduced the LEDGF interaction with labeled AT-probe, whereas competition with unlabeled *IGF2BP2* CpG island and *GAPDH* intron probes had modest effects (Figure 5D). Conversely, LEDGF binding to the CG-probe was disrupted by all three competitors, consistent with lower-affinity, less selective engagement of C/G-rich sequences

**Figure 5.**
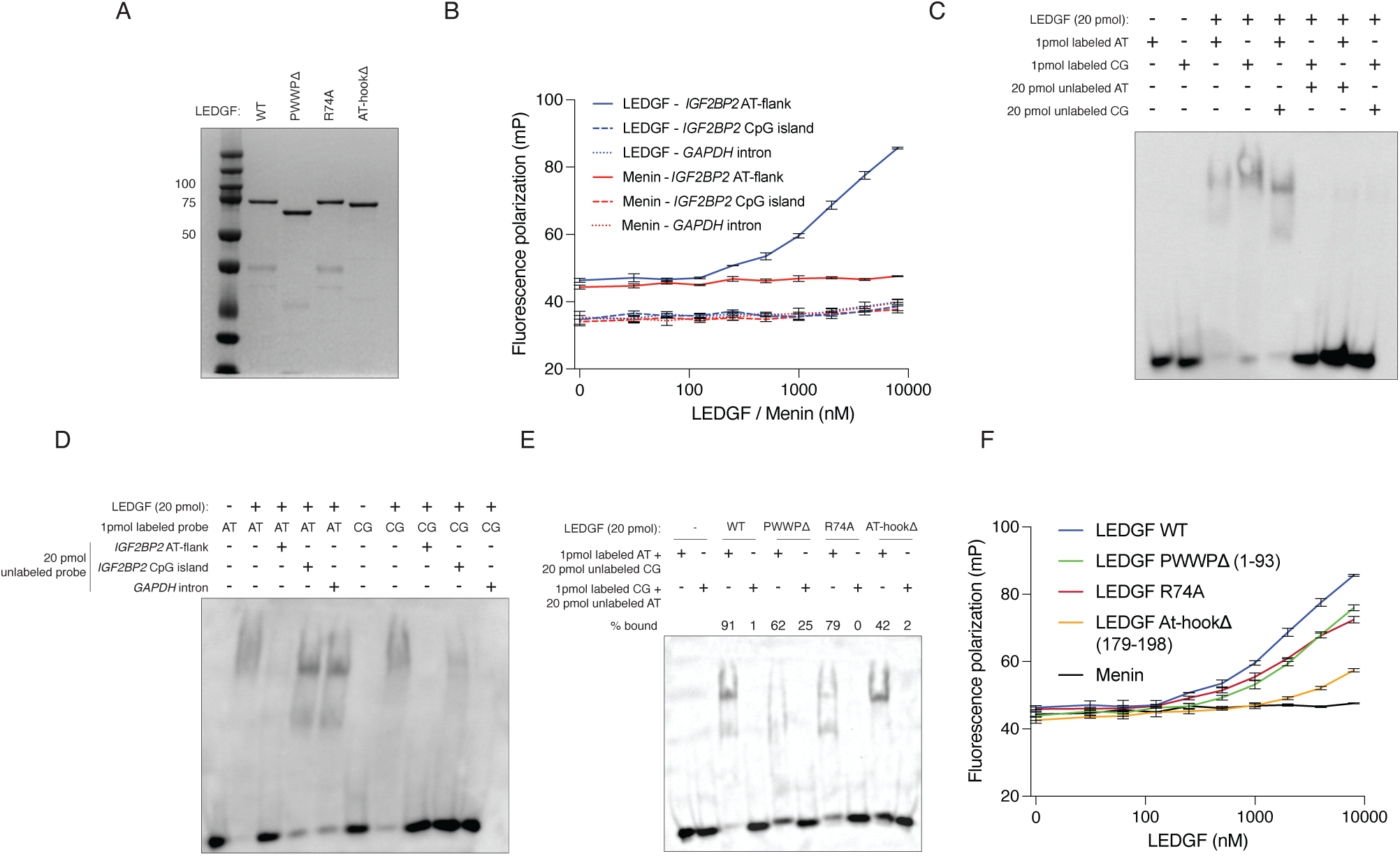
LEDGF preferentially binds to A/T-rich regions. **a,** SDS-PAGE + Coomassie blue staining of 1 μg of affinity-purified 6xHis-LEDGF constructs. **b,** Fluorescence polarization assay (FP) of full-length LEDGF and Menin binding to fluorescein-labeled 25 bp probes derived from an AT-flank and CpG island of the *IGF2BP2* locus and an intronic region from *GAPDH*. **c,** Electrophoretic mobility shift assay (EMSA) of full-length wild-type LEDGF and biotinylated 25 bp AT- and CG-probes. **d,** EMSA of full-length wild-type LEDGF and biotinylated 25 bp AT- and CG-probes with unlabeled competitors. **e,** EMSA with different LEDGF mutant proteins bound to biotin-labeled AT- and CG-probes in the presence of a 20-fold excess unlabeled competitor. The % bound value was calculated using the band intensity of unbound probes. **f,** FP assay measuring the binding of Menin and full-length and mutant LEDGF to the *IGF2BP2* AT-flank probe.

To assess the domains contributing to AT-flank DNA recognition, we measured binding of LEDGF mutants to a biotinylated AT-probe in the presence of excess unlabeled CG-probe. Deletion of either the PWWP or AT-hook domain, as well as the R74A substitution (a PWWP residue previously shown to interact with DNA^54^), reduced LEDGF binding to DNA, yet all mutants retained preferential affinity for the AT-probe (Figure 5E). Fluorescence polarization assays using the *IGF2BP2* AT-flank probe corroborated these results, with the largest reduction observed with the AT-hook deletion (Figure 5F). These findings indicate that multiple regions of LEDGF contribute to engagement of A/T-rich DNA. Collectively, our genomic and biochemical analyses show that LEDGF preferentially binds A/T-rich sequences and may provide a physical link between AT-flanks and the Menin-KMT2A complex.

### LEDGF is required for Menin localization at AT-flanks

To test whether LEDGF is functionally required for Menin-KMT2A engagement of AT-flanked promoters, we used CRISPR/Cas9 to endogenously add an HA-FKBP12 tag to the C-terminus of LEDGF in MV4-11 cells. FKBP12 tagging allows targeted proteasomal degradation upon treatment with the small molecule dTAG13^57^, which tethers the tagged protein to the E3 ubiquitin ligase cereblon. Treatment with dTAG13 resulted in substantial LEDGF degradation after 30 min, becoming undetectable by western blot after 3 hours (Figure S9A). ChIP-seq analysis showed virtually undetectable levels of chromatin-bound LEDGF after 6h of dTAG13 treatment (Figure 6A).

**Figure 6.**
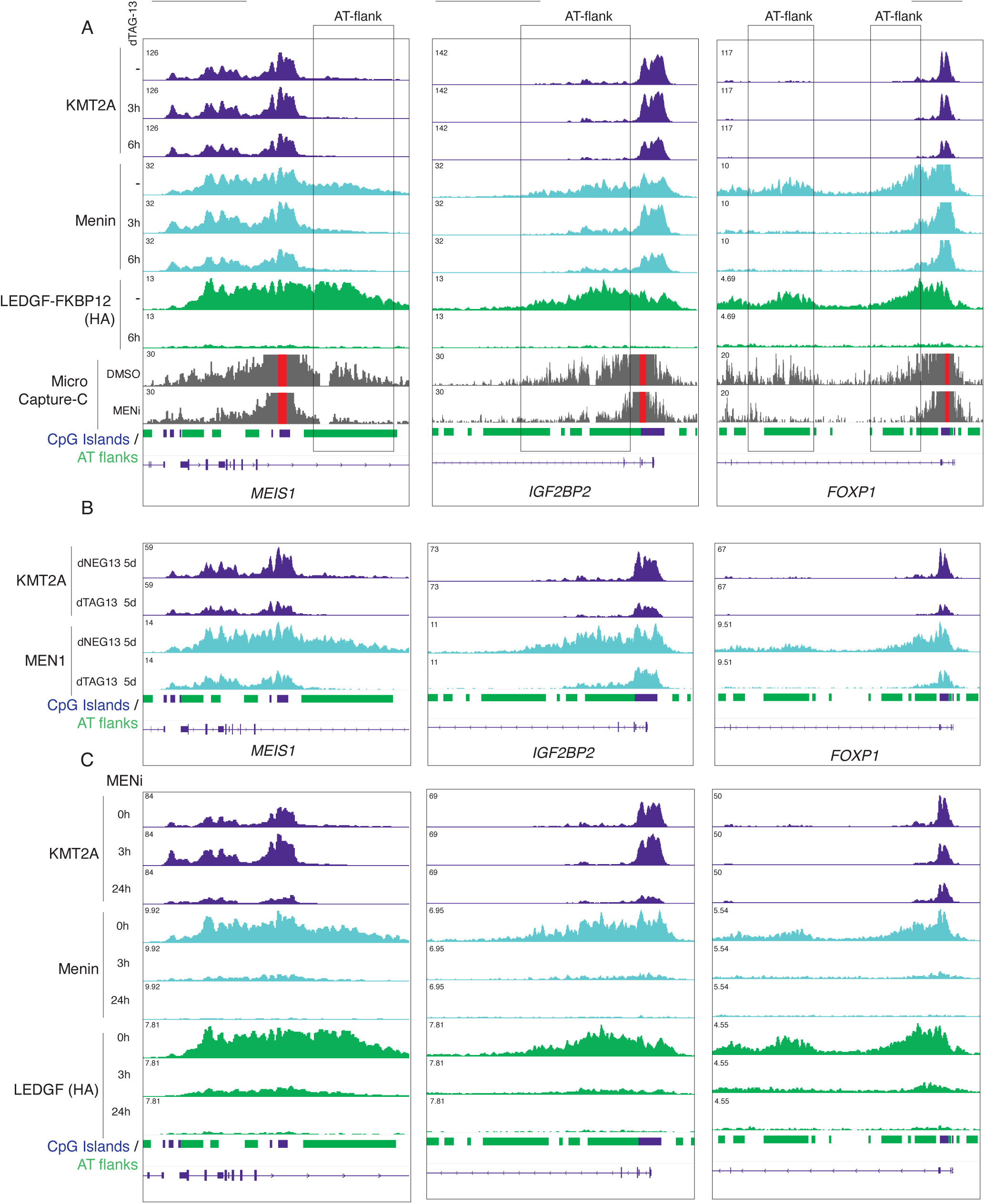
LEDGF is essential for Menin occupancy at AT-flanks. **a,** Overlay of KMT2A, Menin, and LEDGF (HA-tagged) ChIP-seq following acute degradation of LEDGF using the dTAG13 FKBP12 degradation system in MV4-11 cells (500 nM dTAG13), together with Micro Capture-C (MCC) profiles from MV4-11 cells treated with the Menin inhibitor VTP-50469 (MENi; 500 nM, 3 days). Red boxes in the MCC tracks indicate captured regions (CpG islands). **b,** KMT2A and Menin ChIP-seq following 5-day degradation of LEDGF-FKBP12 with dTAG13 (500 nM). The cereblon-engaging molecule dNEG13 (500 nM) was used as a negative control. **c,** ChIP-seq following treatment with Menin inhibitor (MENi: VTP-50469, 500 nM) at the indicated time points. Scale bar represents 10 kb.

ChIP-seq for Menin and KMT2A after 3 h and 6 h of dTAG13 treatment revealed a selective loss of Menin occupancy at AT-flanks, whereas Menin binding at the associated CpG islands remained unchanged (Figure 6A and Figure S9B). KMT2A occupancy at CpG islands was likewise unaffected at these early time points. Notably, the AT-flank regions that lost Menin following LEDGF degradation substantially overlapped regions that engage in long-range contacts with their associated CpG islands, as measured by MCC, and these contacts were disrupted by Menin inhibition (Figures 4 and 6A). Extending LEDGF degradation to 5 days resulted in loss of both Menin and KMT2A from CpG islands genome-wide, whereas occupancy remained unchanged in cells treated in parallel with the cereblon-engaging control compound dNEG13 (Figures 6B and S9B). Prolonged LEDGF degradation also impaired proliferation and increased differentiation and cell death (Figures S9C-D). As observed following Menin inhibition (Figure 2), promoters with greater combined upstream and downstream AT-flank length showed greater loss of KMT2A occupancy after 5 days of LEDGF degradation (Figure S9E). Thus, loss of LEDGF first destabilizes Menin at AT-flanks and is followed by delayed loss of Menin and KMT2A from linked CpG-island promoters.

We next assessed the functional role of the Menin-KMT2A protein-protein interaction on the stability of Menin and LEDGF at AT-flanks. To that end, we performed ChIP-seq for KMT2A, Menin and LEDGF following acute MENi treatment (0, 3, 6 and 24h). MENi rapidly displaced both Menin and LEDGF from AT-flanks and CpG islands, whereas KMT2A remained bound at CpG islands at the earliest time points despite severe loss of Menin and LEDGF (Figure 6C and Figure S9F), and decreased only later (24 h), particularly in genes with large presence of AT-flanks (Figure S9G). Thus, disruption of the Menin-KMT2A interaction rapidly destabilizes Menin and LEDGF occupancy before loss of CpG island-bound KMT2A.

RNA-seq following LEDGF degradation (48h) showed that Menin-KMT2A^high^ targets with more extensive AT-flanks were more strongly downregulated (Figure S9H), mirroring the relationship between AT-flank extent and Menin inhibitor sensitivity described above (Figure 2). Finally, we compared transcriptional changes induced by LEDGF degradation with those induced by Menin inhibition. The two perturbations produced highly correlated expression changes across Menin-KMT2A^high^ genes (Pearson r = 0.796) (Figure S9H), consistent with LEDGF-dependent AT-flank engagement and the Menin-KMT2A interaction acting through a common regulatory mechanism.

Together, these perturbations support a model in which LEDGF-mediated AT-flank recognition and the Menin-KMT2A interaction form an interdependent chromatin-tethering mechanism that sustains KMT2A occupancy at linked CpG-island promoters. Acute LEDGF degradation or disruption of the Menin-KMT2A interaction destabilized the occupancy of the other factor at AT-flanks, indicating that stable engagement of these regions requires both factors.

### The CpG island-AT-flank Composite Promoter Architecture is Associated with Canonical PRC1 Occupancy

Developmental regulators prominent among Menin-KMT2A targets, including HOX factors and MEIS1, are also well-established targets of Polycomb regulation^58,59^. We therefore asked whether the CpG island-AT-flank architecture identified through Menin-KMT2A occupancy might also characterize Polycomb-bound promoters. To address this, we analyzed publicly available ChIP-seq and CUT&RUN datasets for cPRC1 and Polycomb Repressive Complex 2 (PRC2) across multiple cell types^37,60^, including the cPRC1 components BMI1/PCGF4, MEL18/PCGF2, RING1B and CBX proteins, the PRC2 components EZH2 and SUZ12, and the repressive H3K27me3 histone modification catalyzed by PRC2. For each protein or histone modification, genes were divided into groups with high or low promoter occupancy. We restricted the analysis to silent and lowly expressed genes to minimize the confounding effects of the antagonism between active transcriptional machinery and Polycomb occupancy, thereby allowing promoter sequence architecture to be evaluated in a more comparable transcriptional context. Strikingly, the same CNN model trained on Menin-KMT2A occupancy in AML cells distinguished promoters with high cPRC1 occupancy (cPRC1^high^) across cell types from distinct tissues without retraining (ROC AUC > 0.93; Figure 7A). In contrast, classification of PRC2 occupancy and H3K27me3 was weaker and more variable (ROC AUC = 0.72-0.91).

**Figure 7.**
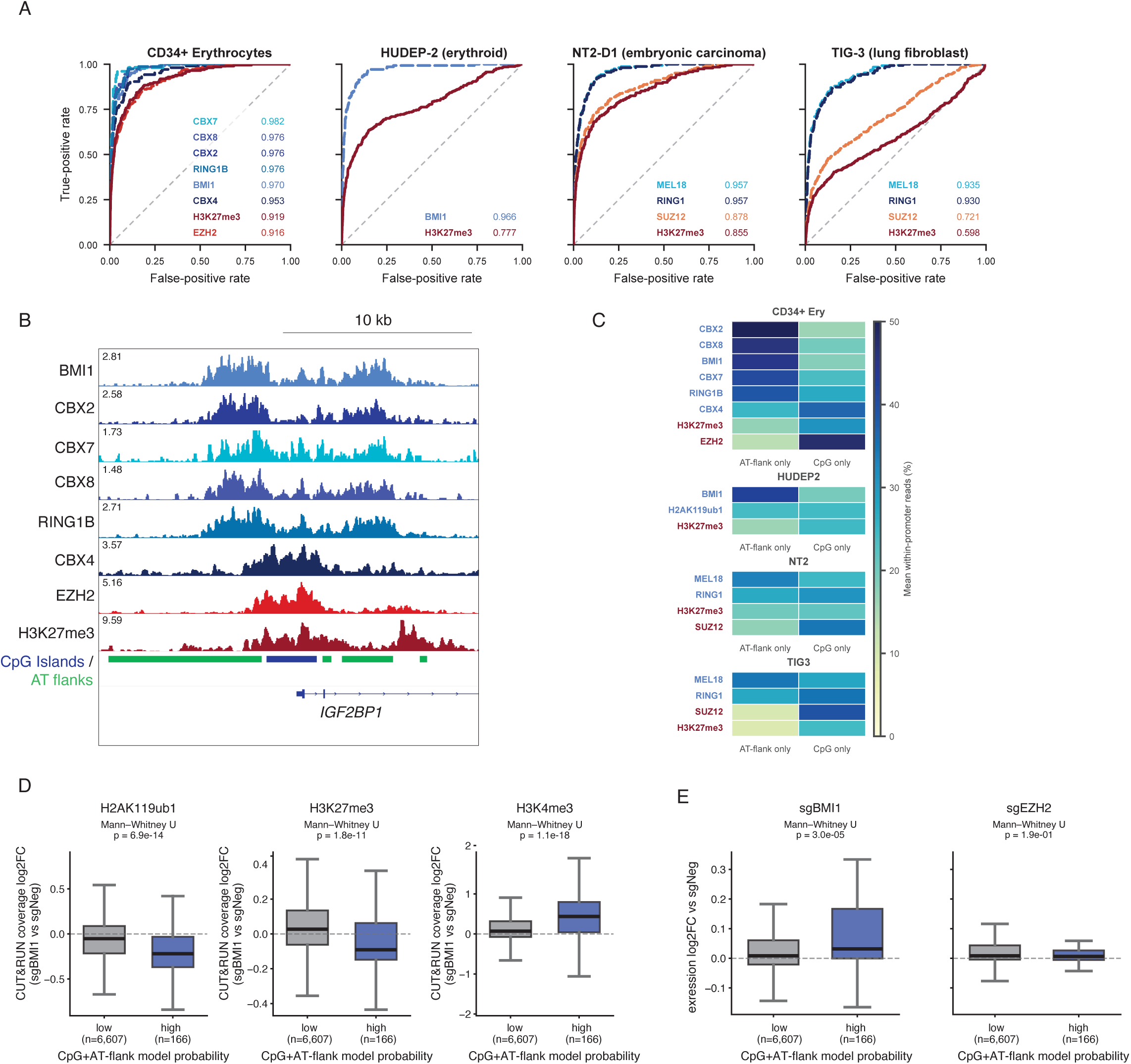
The composite CpG island-AT-flank promoter architecture predicts canonical Polycomb Repressive Complex 1 occupancy across multiple cell types. **a,** Area under the ROC curve for binary classification of high versus low occupancy of cPRC1 components (blue tones) and PRC2 components/H3K27me3 (red tones) at silenced promoters using the CNN model trained on Menin-KMT2A occupancy. b, Representative CUT&RUN profiles of cPRC1/PRC2 components and H3K27me3 at an AT-flanked promoter (I*GF2BP1*) in adult CD34+-derived erythrocytes. c, Average CUT&RUN/ChIP-seq coverage of cPRC1/PRC2 components and H3K27me3 across CpG islands and AT-flanks of their respective target promoters. **d, e,** Changes in CUT&RUN coverage of histone modifications (**d**) and gene expression (**e**) following BMI1 or EZH2 depletion, stratified by CNN model probability, in HUDEP2 cells. Boxes indicate the interquartile range with median; whiskers extend to 1.5× IQR.

Similar to Menin-KMT2A targets (Figure 1D), cPRC1^high^ promoters showed a disproportionate co-occurrence of extensive AT-flanks and CpG islands compared with cPRC1^low^ control promoters (Figure S10A). To determine the relative contributions of A/T- and C/G-rich sequence features to classification, we trained logistic regression classifiers using the average outputs of the corresponding A/T- and C/G-rich filters from the original CNN model as inputs (Figure S10B). C/G-rich features alone classified PRC2 occupancy and H3K27me3 with high accuracy (ROC AUC > 0.93), while inclusion of A/T-rich features produced only a marginal improvement, reflecting the established association of PRC2 with CpG islands^35^. In contrast, classification of cPRC1 occupancy was substantially improved by inclusion of A/T-rich features in addition to C/G-rich features. These sequence differences were reflected in the genomic occupancy patterns of the complexes: BMI1, MEL18, RING1B and CBX2/7/8 were enriched over AT-flanks, whereas EZH2 and SUZ12 were concentrated over CpG islands (Figure 7B-C). CBX4 was an exception, showing greater enrichment over CpG islands despite its association with cPRC1. Furthermore, increased cPRC1 occupancy at AT-flanks was associated with increased PRC2 occupancy at linked CpG islands (Figure S10C).

We next asked whether this architecture was associated with differential dependence on BMI1-containing cPRC1. We reanalyzed published CUT&RUN and RNA-seq datasets from BMI1- and EZH2-depleted HUDEP2 cells^60^, stratifying silenced promoters according to the presence or absence of the CpG island-AT-flank architecture based on CNN model output. Although BMI1 depletion produced only modest global changes in H2AK119ub1 (a repressive histone modification deposited by PRC1) and H3K27me3 levels^60^, promoters containing this architecture showed greater loss of both Polycomb-associated marks, accompanied by a concomitant increase in H3K4me3 (Figure 7D). These promoters were also preferentially derepressed following BMI1 depletion (Figure 7E). In contrast, EZH2 depletion did not preferentially affect expression of the same promoter class. Thus, the CpG island-AT-flank architecture originally identified through Menin-KMT2A occupancy marks a subset of Polycomb-regulated promoters with enhanced dependence on BMI1-containing cPRC1.

## Discussion

A central question in chromatin biology is how regulatory complexes without conventional sequence-specific DNA-binding activity achieve selective genomic localization^1,30,61^. Our findings identify an extended promoter sequence architecture that contributes to this selectivity for the Menin-KMT2A complex. At promoters of a restricted gene set enriched for developmental and cell-fate regulators, CpG islands occur together with large A/T-rich flanking domains that span several kilobases and specify high Menin-KMT2A occupancy. The ability of the same CpG island-AT-flank configuration to distinguish promoters with high cPRC1 occupancy across multiple cell types further indicates that extended promoter sequence composition can contribute to locus-selective chromatin-complex occupancy beyond Menin-KMT2A.

Unlike canonical enhancers, AT-flanks reside in low-accessibility chromatin regions and engage in broad 3D genomic interactions with CpG islands, rather than discrete focal loops^49^. Importantly, genes with high Menin-KMT2A occupancy differ across cell lines, suggesting that AT-flanks alone do not determine recruitment or gene activation. While we show that AT-flanks participate in the maintenance of KMT2A at promoters, the recruitment and activity of Menin-KMT2A may depend on additional regulatory factors that vary with cell state. Accordingly, large intragenic and distal enhancer regions have been found to control the expression of many Menin-KMT2A targets containing AT-flanks in *KMT2A*-r cell lines, indicating a complementary relationship between AT-flanks and enhancers^62,63^. The observation that MENi selectively disrupts AT-flank-CpG island contacts while preserving enhancer-promoter interactions is consistent with potentially distinct roles for these interactions in gene regulation.

Our results also implicate LEDGF as a key reader of AT-flanks. LEDGF has previously been shown to directly bind DNA through an additive mode in which both the PWWP and AT-hook domains affect binding in vitro and in cells^52,64^. Here, we report a previously unappreciated preference of LEDGF for A/T-rich regions, although a similar sequence preference has been observed for the PWWP domain from HRP3, a LEDGF paralog^65^. This preference is retained in mutants lacking either the PWWP or AT-hook domains, suggesting a multi-domain affinity for A/T-rich regions. Our observations, however, do not preclude the existence of other AT-flank “readers” mediating the binding of the Menin-KMT2A complex.

Our findings support a model in which Menin-KMT2A occupancy can be stabilized through two complementary chromatin-tethering mechanisms (Figure S11). Direct association of LEDGF with A/T-rich DNA, coupled through Menin to KMT2A, contributes to the stabilization of KMT2A at the linked CpG-island core. In contrast, KMT2A contains a CXXC domain that directly recognizes unmethylated CpG DNA^32,45^. Consistent with this model, acute disruption of either LEDGF or the Menin-KMT2A interaction initially leaves KMT2A bound at CpG islands, whereas prolonged perturbation leads to its subsequent loss. Moreover, CpG-island-dominated promoters preferentially retain KMT2A following Menin inhibition, while promoters with extensive AT-flanks show greater KMT2A displacement. Thus, the relative contribution of LEDGF-Menin-mediated AT-flank tethering and direct CpG-island recognition by KMT2A may determine the stability and Menin dependence of the complex at individual promoters. This dual-tethering model may help explain why individual target loci exhibit markedly different responses to disruption of the same Menin-KMT2A interaction by clinically approved Menin inhibitors.

The association of this architecture with cPRC1 extends the significance of AT-flanks beyond Menin-KMT2A. This relationship reflected the composite architecture rather than either component alone: cPRC1 proteins preferentially occupied AT-flanks, whereas PRC2 was concentrated at CpG islands. These findings closely parallel the PRC1 tethering elements (PTEs) described by Barrasa et al.^37^, in which cPRC1 occupancy is associated with poly(dA:dT)-rich sequence features that are frequently juxtaposed to CpG-rich regions favoring PRC2 binding. Barrasa et al. proposed that distinct sequence determinants for PRC1 and PRC2 combine in different proportions across Polycomb targets. Our results provide a predictive framework for this sequence organization, identifying A/T- and C/G-rich features carrying sufficient information to distinguish cPRC1^high^ promoters from cPRC1^low^ promoters across multiple cell types. Remarkably, this predictive architecture was learned entirely from Menin-KMT2A occupancy, without incorporating cPRC1 occupancy information, revealing a shared promoter architecture that can distinguish both repressed cPRC1^high^ and active Menin-KMT2A^high^ promoters. The convergence of activating and repressive complexes on a shared sequence architecture is reminiscent of the dual occupancy of PcG and TrxG response elements in *Drosophila*, although whether these represent functionally analogous elements remains to be established.

Together, our findings suggest that the CpG island-AT-flank configuration may represent a sequence-encoded regulatory platform enriched at developmental genes, whose functional output depends on the chromatin complexes that engage it. Understanding how cellular context determines whether this shared architecture is engaged by active Menin-KMT2A or repressive Polycomb complexes may reveal a broader principle governing the regulation of developmental gene programs.

### Limitations of the study

The function of AT-flanks was established through endogenous deletions and other perturbations rather than ectopic insertion or sequence substitution, precluding direct assessment of whether AT-flanks are sufficient to recruit LEDGF and Menin. The large sizes of the AT-flanks and CpG islands associated with high Menin-KMT2A occupancy pose a substantial technical challenge for manipulating these genomic elements. Furthermore, the association between the composite CpG island-AT-flank architecture and cPRC1 occupancy remains correlative and has not been tested by direct perturbation of the underlying genomic elements or identification of the molecular mechanism that recognizes AT-flanks.

## Data and code availability

Processed and raw ChIP-seq, RNA-seq and MCC data have been deposited to GEO and will be made available upon publication. The computational analyses in this study were written in C and Python and are available at: https://github.com/armstrong-lab/AT-flanks

## Acknowledgments

We thank all members of the Armstrong laboratory for helpful discussions. P.R.F. is supported by a Momentum Fellowship from The Mark Foundation for Cancer Research. S.A.A. was supported by the St. Jude Children’s Research Hospital Collaborative research program.

## Author contributions

P.R.F. conceived the project, designed and performed experiments, and conducted computational analyses. S.A.A. supervised the project and contributed to project conception and experimental design. B.R. and E.T.E. performed experiments. R.T. and G.W. generated the MV4-11 LEDGF-FKBP12 cell line. T.Z.B. performed logistic regression analysis of DNA k-mers. T.Z.B., J.B., and R.A.I. provided guidance on computational and statistical analyses. V.S. and T.A.M. provided guidance on MCC protocols and data analysis. T.A.M. and J.A.C. provided overall scientific guidance. P.R.F. and S.A.A. wrote the manuscript.

## Methods

### Cell Culture, Drug Treatment and Virus production

MV4-11 and OCI-AML3 cells were cultured in RPMI medium. HeLa and 293T cells were cultured in DMEM medium. 10% fetal bovine serum, antibiotics and L-glutamine were supplemented in all cell cultures. The Menin inhibitor VTP-50469, SNDX-5613 (Revumenib), dTAG13 and dNEG13 (dTAG-13-NEG) were obtained from MedchemExpress. RNA Poll initiation inhibitor triptolide was obtained from Selleck. Viral supernatants were generated by transfection of 293T cells with 3^rd^ generation lentivirus vectors using the X-tremeGene transfection reagent (Millipore). Transfection of MV4-11 cells was done via spinoculation of undiluted viral supernatant at 950g for 90 min at 37°C. Fresh medium was added immediately after centrifugation. For HeLa cells, the viral supernatant was added to culture at a 1:4 ratio overnight.

### CD34+ Primary Cells

Umbilical cord blood was obtained at the Pasquarello Tissue Bank at Dana-Farber Cancer Institute. Peripheral blood mononuclear cells were extracted using Ficoll-paque (Cytiva). CD34+ cells were purified utilizing the EasySep Human Cord Blood CD34 Positive Selection Kit II (Stemcell Technologies) and flash frozen in liquid nitrogen.

### ChIP-sequencing

Twenty million cells (or 2 million frozen CD34+ cord blood cells) were fixed with 2 mM disuccinimidyl glutarate for 30 min at room temperature followed by a 10 min fixation with 1% formaldehyde. Fixation was quenched with 100 mM Tris pH 7.5 and 125 mM glycine. For CD34+ cord blood samples and short term LEDGF-FKBP12 /MENi treatments samples (Figure 6), only formaldehyde fixation was used. For KMT2A, Menin and HA ChIPs, chromatin was washed once with SDS buffer (100 mM NaCl, 50 mM Tris pH 8.0, 5mM EDTA pH 8, 0.5% SDS) and resuspended in SDS buffer + Triton dilution buffer (100 mM Tris pH 8, 100 mM NaCl, 5 mM EDTA pH 8, 5% Triton X-100) at a 1:3 ratio prior to sonication. For all other ChIPs, chromatin was washed with LB1 (50 mM HEPES pH 7.4, 140 mM NaCl, 1 mM EDTA, 10% glycerol, 0.5% IGEPAL CA-630, 0.25% triton X-100), LB2 (10 mM Tris pH 8, 200 mM NaCl, 1 mM EDTA, 0.5 mM EGTA) and resuspended in LB3 (10 mM Tris pH 8, 100 mM NaCl, 0.1% sodium deoxycholate, 0.5% sodium lauroylsarcosine, 1% Triton X-100, 1mM EDTA, 0.5 mM EGTA) before sonication. Protease inhibitors were added to all buffers. Chromatin was sonicated to an average size of 200-600 bp using a Covaris E220 sonicator followed by centrifugation for 20 min at 20,000g at 4 °C. All samples were incubated overnight at 4°C with antibodies (KMT2A: Bethyl, A300-086A, 5 μg; Menin: Bethyl, A300-105A, 5 μg; LEDGF: Bethyl, A300-848A, 1 μg; HCF-1: Bethyl, A301-400A, 5 μg; ENL: Millipore, ABE2596, 5 μg, RNA Pol II: Abcam, ab26721, 5 μg; H3K9ac: Abcam, ab4441, 5 μg; H3K27ac: Diagenode, C15410196, 5 μg; H3K4me3: Abcam, ab8550, 5 μg, HA: Abcam, ab9110, 5 μg) pre-coupled with protein A (rabbit antibodies) or protein G (mouse antibodies) Dynabeads (Invitrogen). KMT2A, Menin and HA ChIP samples were diluted 4-fold in ChIP Dilution Buffer (16.7 mM Tris pH 8, 167 mM NaCl, 1.1% Triton X-100, 1.1 mM EDTA). For all ChIPs, beads were washed twice for 5 min in Low Salt Buffer (50 mM Tris pH 8, 150 mM NaCl, 0.1% SDS, 0.5% sodium deoxycholate, 1% Triton X-100, 1mM EDTA), High Salt Buffer (50 mM Tris pH 8, 500 mM NaCl, 0.1% SDS, 0.5% sodium deoxycholate,1% Triton X-100, 1mM EDTA), LiCl buffer (50 mM Tris pH 8, 250 mM LiCl, 1% sodium deoxycholate, 1% IGEPAL CA-630 1mM EDTA). Beads were then quickly washed twice with TE buffer (10 mM Tris pH 8, 1mM EDTA) without removal from the magnet and resuspended in SDS elution buffer (50 mM Tris pH8, 200 mM NaCl, 1% SDS, 1mM EDTA) followed by overnight incubation at 65°C. DNA was purified using AMPure XP beads (Beckman Coulter) at a 1:1 ratio and eluted in TE buffer. For RNA Pol II samples, beads were washed and eluted in a similar manner with the addition of two washes of Mixed Micelle Buffer (20 mM Tris pH 8, 150 mM NaCl, 5mM EDTA, 5.2% sucrose, 0.2% SDS, 1% Triton X-100) prior to Low Salt Buffer washes. ChIP-seq libraries were prepared using the ThruPLEX DNA-Seq kit (Takara) and sequenced using an Illumina NextSeq 550 sequencer, generating 37 bp paired-ended reads.

### ChIP-sequencing analysis

Reads were aligned to the human genome (hg38) using STAR 2.7.5 and duplicates were removed using Picard tools v2.9.4. Promoter coverage was calculated using bedtools (v 2.31.1). For calculating promoter coverage, promoters in which the input counts were higher than 3 standard deviation from the trimmed mean counts (>2.5 and < 97.5 percentiles) were eliminated. Bigwig files for genome coverage visualization and average promoter binding profiles were calculated using DeepTools (v3.5.6) using only primary alignments. On Figures 2,3F, 6 and 7B, genome coverage was normalized by the Reads Per Million method. On Figure 3 (A,C), genome coverage was normalized by the mean Menin read count per kb in the promoter region (-3 to +20 kb) among Menin-KMT2A^high^ targets with the exception of *PBX3* and *CDK6*, which were disrupted by CRISPR/Cas9 deletions. On Figure 3D,E, KMT2A promoter read counts from WT and sgMEN1 HeLa cells were normalized using the median ratio method in a similar manner to DESeq2^66^. On Figure S9B, E, F and G, FPKM normalization was used to account for the different feature sizes.

### AT-flank conservation and enhancer comparison

AT-flanks were defined as stretches of DNA with a frequency of AAA or TTT higher than 0.1 in a 500 bp window. Adjacent flanks less than 500 bp apart were merged. CpG islands definitions were downloaded from the UCSC Genome Browser. Apart from the Bigwig files generated by our study, the following Bigwig files were downloaded from the CistromeDB database^67^: BRD4 (CistromeDB number: 93576^68^) and CDK8 (CistromeDB number: 56375^41^). MV4-11 ATAC-seq data Bigwig files were downloaded from ChIP-Atlas^69^ (accession number: SRX6865324^70^). The mammalian conservation (PhastCons score from 470 mammal assemblies) and ENCODE Candidate Cis-Regulatory Elements (cCREs)^43^ were downloaded from the UCSC Genome Browser. The enhancer regions in MV4-11 were defined in a previous publication^71^ using the activity by contact (ABC) model^72^.

### Western Blot

Western blot was done using the direct lysis of 100,000 cells into 100 μl of 1x LDS buffer (Thermo Fischer Scientific). NuPAGE Bis-Tris gels were used in MOPS buffer, and membrane transfer was performed with an iBlot3 machine (Thermo Fischer Scientific). The membrane was blocked in 5% milk and incubated overnight with anti-HA (Abcam, ab9110, 1:1000 dilution) followed by HRP-conjugated anti-rabbit IgG secondary antibody (Cell Signaling Technologies, #7074, 1:2000 dilution). Membrane was imaged using Millipore Immobilon ECL substrate and a ChemiDoc MP imager (BioRad). For loading control, the membrane was stripped with Restore buffer (Thermo Fischer Scientific) and reblotted with an anti-Beta-Actin-HRP antibody (Cell Signaling Technologies, 5125, 1:1000 dilution).

### RT-qPCR

RNA was extracted with the RNeasy kit from Qiagen and cDNA was generated using the High-Capacity cDNA Reverse Transcription Kit (Invitrogen). *PBX3* and *CDK6* gene expression was measured by Taqman Gene Expression assays using a QuantStudio6 Flex machine (*PBX3*: Hs00608415_m1, *CDK6:* Hs01026371_m1 and *GAPDH*: Hs02786624_g1).

### Flow cytometry, Differentiation and Proliferation Assays

The myeloid differentiation of LEDGF-FKBP12 MV4-11 cells was measured via flow cytometry using the PE-anti-human CD14 (Biolegend antibody, 1:400 dilution). Apoptosis was measured using the APC Annexin V Apoptosis Detection Kit (BD Biosciences). Flow cytometry was performed in a LSRFortessa cytometer (BD Biosciences). Cell proliferation was measured manually with a hemocytometer.

### RNA-seq

For the analysis of the transcriptional effects of SNDX-5613 in MV4-11 cells, public available fastq files were downloaded from Gene Expression Omnibus (GEO; GSE196035)^73^. Paired-ended reads (paired-end 37-mers) were aligned to the human genome using STAR (v2.7.5a), sorted and duplicates marked/removed with picard pipeline tools (v2.9.4). Raw per-gene counts were calculated with HTSeq (htseq-count, v0.6.1pl). Differential RNA-seq expression was calculated using the BioConductor DESeq2 package (v1.24,0), using raw unnormalized per-gene counts from deduplicated BAMs.

For the RNA-seq analyses after short term LEDGF degradation and Menin inhibition with SNDX-5613, 100,000 cells were resuspended with DNA/RNA-shield (Zymo) and delivered to Plasmidsaurus for RNA-seq and analysis. Gene-level quantification and differential expression analysis were performed using the Plasmidsaurus RNA-Seq pipeline (demultiplexing, fastp quality filtering, STAR alignment to hg38, UMI-based deduplication, featureCounts quantification, TMM normalization, edgePython^74^ differential expression).

### Convolutional Neural Network

To define the Menin-KMT2A^high^ and Menin^low^ categories, negative binomial distributions were fitted to promoter read counts of KMT2A and Menin in each cell line. Only expressed genes were considered, defined by TPM > 2 from the expression dataset downloaded from the DepMap project on January 2025. Menin-KMT2A^high^ set was defined as expressed promoters with Menin occupancy negative binomial CDF > 0.99995 and KMT2A CDF > 0.995. The Menin^low^ set consisted of expressed genes with Menin occupancy negative binomial CDF < 0.8. The final training set consisted of the union of Menin-KMT2A^high^ and Menin^low^ sets in MV4-11 and OCI-AML3 cell lines. 10% of the training set was split into a validation set to determine training stop. An external testing set consisted of the Menin-KMT2A^high^ and Menin^low^ promoters defined in Hela cells. Promoters that were also present in the training and validating sets were eliminated from the test set. As CpG islands are essential for KMT2A binding^32^, promoters without CpG islands were discarded. Given the high imbalance of the training data (43 Menin-KMT2A^high^ vs. 7910 Menin^low^), the Menin-KMT2A^high^ were oversampled to force a 1:1 balance between classes. Menin and KMT2A ChIP-seq for MV4-11 and HeLa were performed as described above. The following fastq files were downloaded from Sequence Read Archive and processed as described above: OCI-AML3 Menin (SRX14127512, SRX14127566), OCI-AML3 KMT2A (SRX18455080, SRX18455076)^23^ and MOLM13 Menin (SRX10269611, SRX10269609)^71^.

The convolutional neural network consisted of two convolution layers, the first with 10 × 5 bp kernels, 5-wide max pooling and ReLU activation function. The second layer had 128 × 10 kernels with 4950-wide average pooling and ReLU activation function. The output of the second layer was flattened and summarized into a single value through a linear layer with a dropout ratio of 0.2 and a final sigmoid activation function. The code was implemented using pytorch and executed in a T4 high-RAM GPU using Google Colab. The integrated gradients attribution analysis on input nucleotides and kernel layers were done using the Captum library.

### Logistic Regression

For each combination of k-mer length and genomic interval, an elastic net-regularized logistic regression model was trained to classify promoters as high or low binding, based on the previously defined ChIP-seq derived labels. The model used a balanced combination of L1 and L2 penalties with a regularization strength of C=0.1 to identify a sparse set of predictive k-mers while minimizing overfitting. Model performance was assessed using a 5-fold stratified cross-validation scheme, with the Area Under the Receiver Operating Characteristic Curve (AUC) as the primary metric.

### CRISPR/Cas9 AT-flank Deletions

For deleting AT-flanks in MV4-11 cells, a stable transduction of Cas9 was first performed using lentiviral vectors. Next, sgRNAs were transduced and selected with puromycin targeting the borders of the targeted AT-flanks. A total of 4 sgRNAs per cut site (8 per AT-flank) were simultaneously transduced. Single cell clones were derived via limiting dilution and verified via PCR. The targeting of Menin in HeLa cells followed a similar procedure but using only a single sgRNA targeting Menin. Single cell clones were also derived via limiting dilution. sgRNA sequences are described on Table S2.

### LEDGF-FKBP12-HA Cell Line Generation and Characterization

Endogenous insertion of the FKBP12-2xHA-p2A-mNeonGreen construct into the 3’end of the *PSIP1* locus (which encodes LEDGF) in MV4-11 cells was performed by electroporation. Briefly, cells were resuspended in SG Nucleofector Solution supplemented with the appropriate supplement (Lonza) and mixed with the Cas9/sgRNA RNP complex and Alt-R™ Cas9 Electroporation Enhancer (Integrated DNA Technologies). The RNP complex was delivered using the 4D-Nucleofector™ system (Lonza). The culture medium was replaced the following day. mNeonGreen-positive cells were sorted, followed by single-cell cloning to establish homozygous edited cell lines. sgRNA sequence: tgaaatcgcgagtcatgttt.

### PRC1/2 Occupancy Prediction

CUT&RUN (CD34+ Erythrocytes and HUDEP2) and ChIP-seq (NT2 and TIG3) fastq files were downloaded from the GEO database (Table S3). ChIP-seq samples were analyzed as described above. CUT&RUN samples (paired-end reads) were aligned using Bowtie2 in very-sensitive local mode (--local --very-sensitive-local), retaining only concordant pairs with insert sizes of 10–700 bp (--no-mixed --no-discordant -I 10 -X 700). Promoter coverage was calculated using the -11.5 to +11.5 kb window around the TSS, given the greater centrality of PRC occupancy compared to Menin-KMT2A complex, which used the -3 to +20 kb window.

Positive and negative PRC target sets were defined at the promoter level using chromatin occupancy together with matched gene expression. Because PRC occupancy is strongly anti-correlated with expression, we used the promoter coverage at expressed genes to estimate the background ChIP-seq signal at PRC1/2-negative promoters. For each Polycomb protein or histone mark, expressed genes (log2(TPM + 1) > 0.5) were used to fit a negative-binomial distribution to the promoter coverage counts. The fitted 80th and 99.9999th percentiles defined the lower and upper occupancy thresholds, respectively. Labels were assigned only among low-expression promoters, defined as log2(TPM + 1) < 0.5. Matched expression came from GSE218232 for CD34 and HUDEP2 and from DepMap-CCLE for NT2 and TIG3. For HUDEP2 histone marks, only sgNegative samples were used. Biological replicates were normalized using DESeq2-style median-of-ratios size factors and then averaged. Perturbed HUDEP2 samples were not used to construct the CNN target labels. Replicate 1 was retained for the principal cross-cell-line ROC and precision–recall figures when separate replicate labels existed. These PRC labels were used to evaluate the pretrained CNN’s continuous promoter probabilities; the later F1-optimized probability cutoff was used only for downstream model-group comparisons, not to establish the experimental positive and negative sets. For figure 7E, processed gene-level RSEM outputs for HUDEP2 cells were obtained from GSE218232^60^ and analyzed with tximport and DESeq2. The analysis was restricted to the experimentally defined HUDEP2 BMI1 target sets and divided using the F1-optimized CNN probability threshold of 0.846. To assess whether AT- and CpG-associated sequence features provided complementary information for Polycomb target classification, we fitted class-weighted logistic-regression models to experimentally defined positive and negative promoters. Models containing the mean standardized activation of CpG-associated CNN filters alone were compared with models containing both CpG- and AT-associated filter activations. Performance was evaluated by repeated stratified fivefold cross-validation with three repeats and summarized using held-out ROC AUC. An analogous analysis was performed using literal CpG-island and AT-flank overlap lengths within the -11.5 to +11.5 kb promoter window.

### Micro Capture-C

The Micro Capture-C protocol was performed according to Hamley et al. 2023^48^. Briefly, 30 million MV4-11 cells treated with DMSO (3 days), VTP-50469 (500 nM, 3 days) or triptolide (500 nM, 1 h) were fixed with 2% formaldehyde for 10 minutes and permeabilized with 1% digitonin. The chromatin was then fragmented using MNase and ligated overnight with T4 ligase. DNA was purified with DNeasy columns (Qiagen) and sequencing adaptors were added with the NEBNext Ultra II kit (New England Biolabs). Finally, target regions were captured using the KAPA HyperCapture Reagent kit (Roche) and sequenced using 150 bp paired-end reads. The biotinylated oligos used for capture are described on Table S2. All regions were captured with one oligo, with the exception of the AT-flank of *MEIS1*, in which two oligos were used. The data analysis was performed using the PERL scripts made available by the authors.

### Protein expression and purification

Full length Menin was a gift from Dr. Eric Fischer’s Lab, and it was purified from *E. coli* as previously described^73^. Full length LEDGF was cloned into a pET28 vector with a N-terminal 6x His tag. LEDGF mutant constructs were generated using the Q5 site-directed mutagenesis (New England Biolabs). The vectors were transformed into Rosetta 2(DE3) *E. coli* (Novagen) and expression was induced in 3 liters of bacterial culture with 0.5 mM IPTG overnight at 16°C. The bacterial pellet was lysed in Bacterial Lysis Buffer (10 mM HEPES pH 7.4, 300 mM NaCl, 10% glycerol, 10 mM imidazole, 5 mM beta-mercaptoethanol, 1mM PMSF, 1500U benzonase, protease inhibitors), sonicated with a probe tip sonicator (2 min total time, 10s ON / 20s OFF, 40% amplitude) and centrifuged at 108,000g for 1h at 4°C. The supernatant was applied to 3 ml of TALON Metal affinity resin (Takara) and incubated for 1h at 4°C with rotation. The resin was transferred to a gravity column and washed with 20 volumes of lysis buffer, 10 volumes of High Salt Wash (10 mM HEPES pH 7.4, 1M NaCl, 10% glycerol, 10 mM imidazole, 5 mM beta-mercaptoethanol) followed by 10 volumes of lysis buffer. The protein was recovered with 3 x 10ml elution with Elution Buffer (20 mM HEPES pH 7.4, 300 mM NaCl, 10% glycerol, 200 mM imidazole, 5mM beta-mercaptoethanol). Sample was concentrated using the Amicon Ultra-15 Centrifugal filter (100 kDa MWCO, Millipore) and dialyzed overnight in Dialysis Buffer (20 mM HEPES pH 7.4, 300 mM NaCl, 10% glycerol, 5mM beta-mercaptoethanol) at 4°C. Samples were aliquoted, flash frozen and stored at -80°C.

### Electrophoretic Mobility Shift Assay (EMSA)

20 pmol of recombinant LEDGF was incubated in EMSA buffer (25 mM Tris pH 7.4, 5 mM MgCl_2_, 120 mM NaCl, 10% glycerol, 2 mM DTT and 50 μg/ml BSA) with 1pmol of biotin-labeled 25 bp DNA probe with or without 20 pmol of unlabeled probes for 15 min at room temperature. The total sample volume was 20 μl. Samples were separated by a native 6% 0.5X TBE gel and transferred to a nylon membrane via wet transfer for 1h at 30V in 0.5x TBE buffer. The DNA was crosslinked to the membranes with a UV illuminator (120mJ/cm^2^, Stratagene) and blocked with 4% BSA in PBS for 1h at room temperature. The membranes were then incubated with Streptavidin-HRP (abcam ab7403, 1:40,000 dilution) for 30 min in blocking buffer, washed with PBS for 5 min, APS buffer (10% fetal bovine serum, 1% Triton X-100 in PBS) for 30 min and again 5 min PBS. The membranes were treated with Immobilon Chemiluminescent Substrate (Millipore) for 5 min and imaged.

### Fluorescence Polarization Assay

Varying amounts (0 to 10 μM) of recombinant Menin or LEDGF were incubated with 20 nM of fluorescein-labeled 25 bp DNA probes in EMSA buffer for 15 minutes at room temperature. Fluorescence polarization was measured using a CLARIOstar Plus plate reader (BMG Labtech).

