## Supplemental Figures for "A Unique Promoter Architecture Defines Menin-KMT2A Complex Selectivity"

### Supplemental Figure1

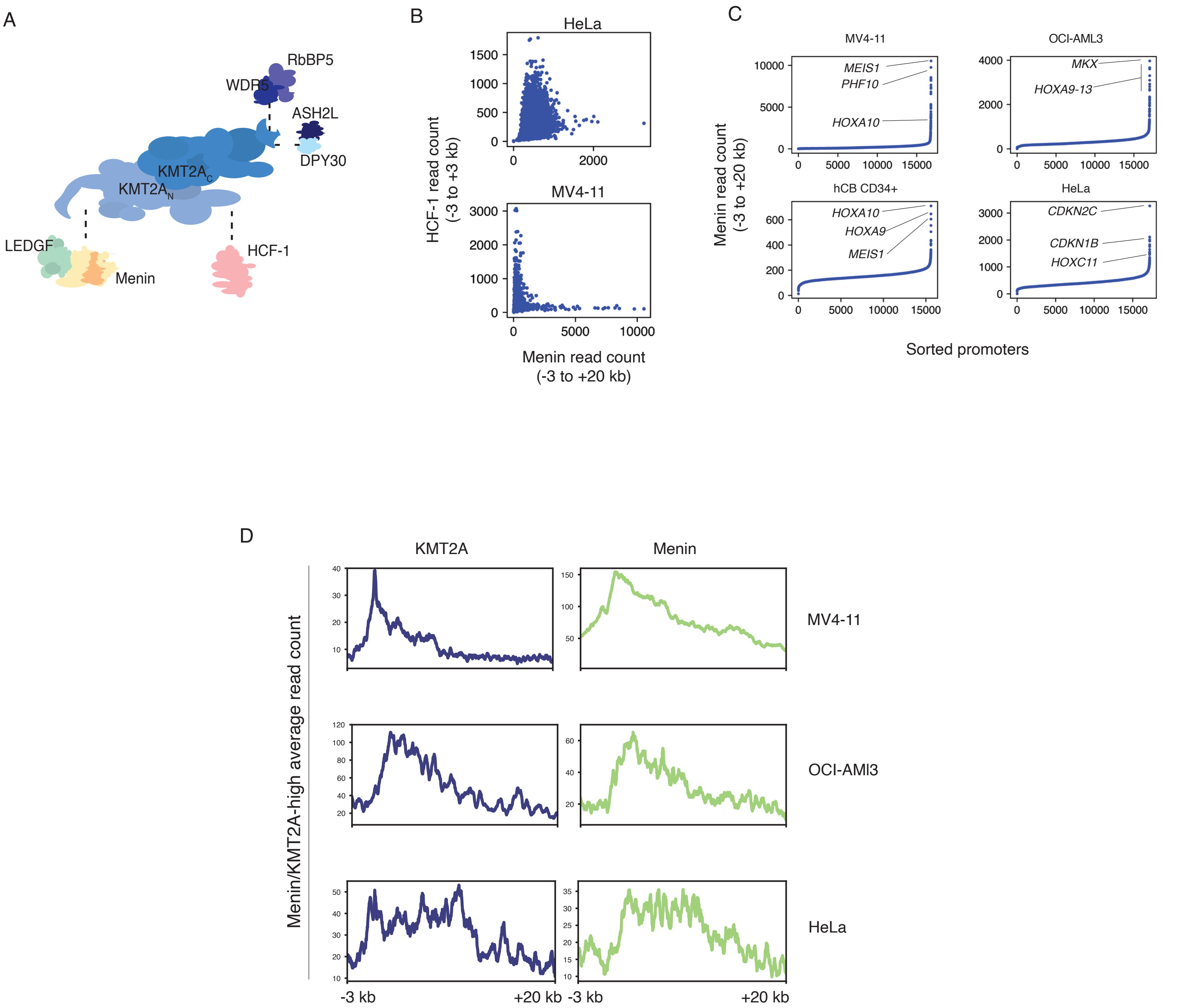


**Figure S1. Promoter occupancy of KMT2A complex members HCF-1 and Menin. a,** Schematic view of the KMT2A complex. **b,** Scatter plot comparing Menin and HCF-1 ChIP-seq read counts in MV4-11 and HeLa cells. **c,** Ranked Menin occupancy (ChIP-seq) in MV4-11 (KMT2A-r), OCI-AML3 (NPM1c-mutant), human cord blood CD34+ cells (normal hematopoietic progenitors) and HeLa (cervical carcinoma). **d,** ChIP-seq profiles of KMT2A and Menin at Menin-KMT2A$^{high}$ promoters (n = 43) in MV4-11, OCI-AML3 and HeLa cells.

### Supplemental Figure 2

A

HOMER known motifs

| Rank | Motif | Name | P-value | log P-pvalue | q-value (Benjamini) | # Target Sequences with Motif | % of Targets Sequences with Motif | # Background Sequences with Motif | % of Background Sequences with Motif |
|---|---|---|---|---|---|---|---|---|---|
| 1 | 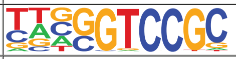 | HINFP(Zf)/K562-HINFP.eGFP-ChIP-Seq(Encode)/Homer | 1e-4 | -1.037e+01 | 0.0148 | 44.0 | 100.00% | 6689.3 | 79.00% |
| 2 | 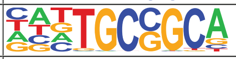 | Zfp57(Zf)/H1-ZFP57.HA-ChIP-Seq(GSE115387)/Homer | 1e-3 | -6.967e+00 | 0.2223 | 44.0 | 100.00% | 7227.8 | 85.36% |
| 3 | 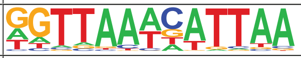 | Hnf1(Homeobox)/Liver-Foxa2-Chip-Seq(GSE25694)/Homer | 1e-2 | -5.603e+00 | 0.5799 | 41.0 | 93.18% | 6476.9 | 76.49% |
| 4 | 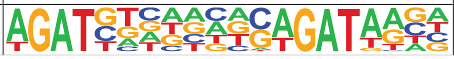 | GATA3(Zf),DR8/iTreg-Gata3-ChIP-Seq(GSE20898)/Homer | 1e-2 | -4.853e+00 | 0.9212 | 30.0 | 68.18% | 4145.3 | 48.95% |

HOMER *de novo* motifs

| Rank | Motif | P-value | log P-pvalue | % of Targets | % of Background | STD(Bg STD) |
|---|---|---|---|---|---|---|
| 1 | 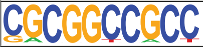 | 1e-16 | -3.762e+01 | 81.82% | 21.89% | 5376.0bp (5895.9bp) |
| 2 | 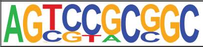 | 1e-14 | -3.391e+01 | 65.91% | 13.55% | 5020.9bp (6173.4bp) |
| 3 | 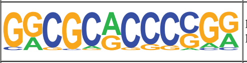 | 1e-14 | -3.357e+01 | 61.36% | 11.31% | 5860.1bp (5890.4bp) |
| 4 | 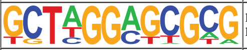 | 1e-14 | -3.354e+01 | 45.45% | 4.83% | 6241.5bp (6413.8bp) |
| 5 | 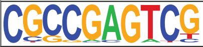 | 1e-14 | -3.346e+01 | 59.09% | 10.24% | 5456.8bp (5896.0bp) |
| 6 | 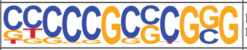 | 1e-12 | -2.920e+01 | 70.45% | 19.17% | 4539.8bp (5407.2bp) |
| 7 | 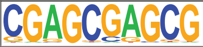 | 1e-12 | -2.825e+01 | 50.00% | 8.28% | 4908.4bp (5362.7bp) |

B

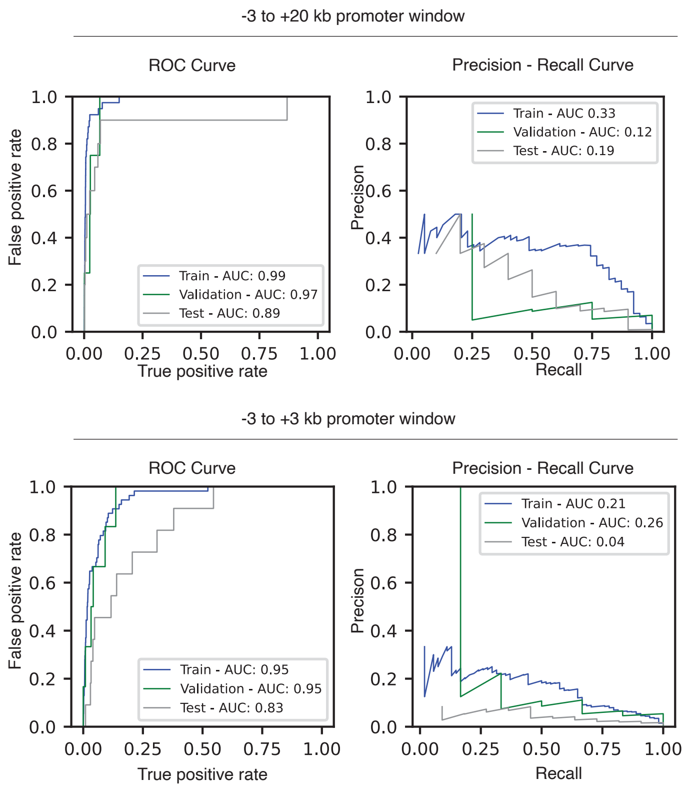


**Figure S2. Large promoter input windows improve CNN model performance. a,** HOMER motif enrichment analysis comparing Menin-KMT2A$^{high}$ promoters and Menin$^{low}$ promoters within a -3 to +20 kb window. Known and de novo motif outputs are shown. **b,** ROC and precision-recall curves comparing CNN model performance using -3 to +3 kb and -3 to +20 kb input windows. ROC, Receiver operating characteristic; AUC, area under the curve.

### Supplemental Figure 3

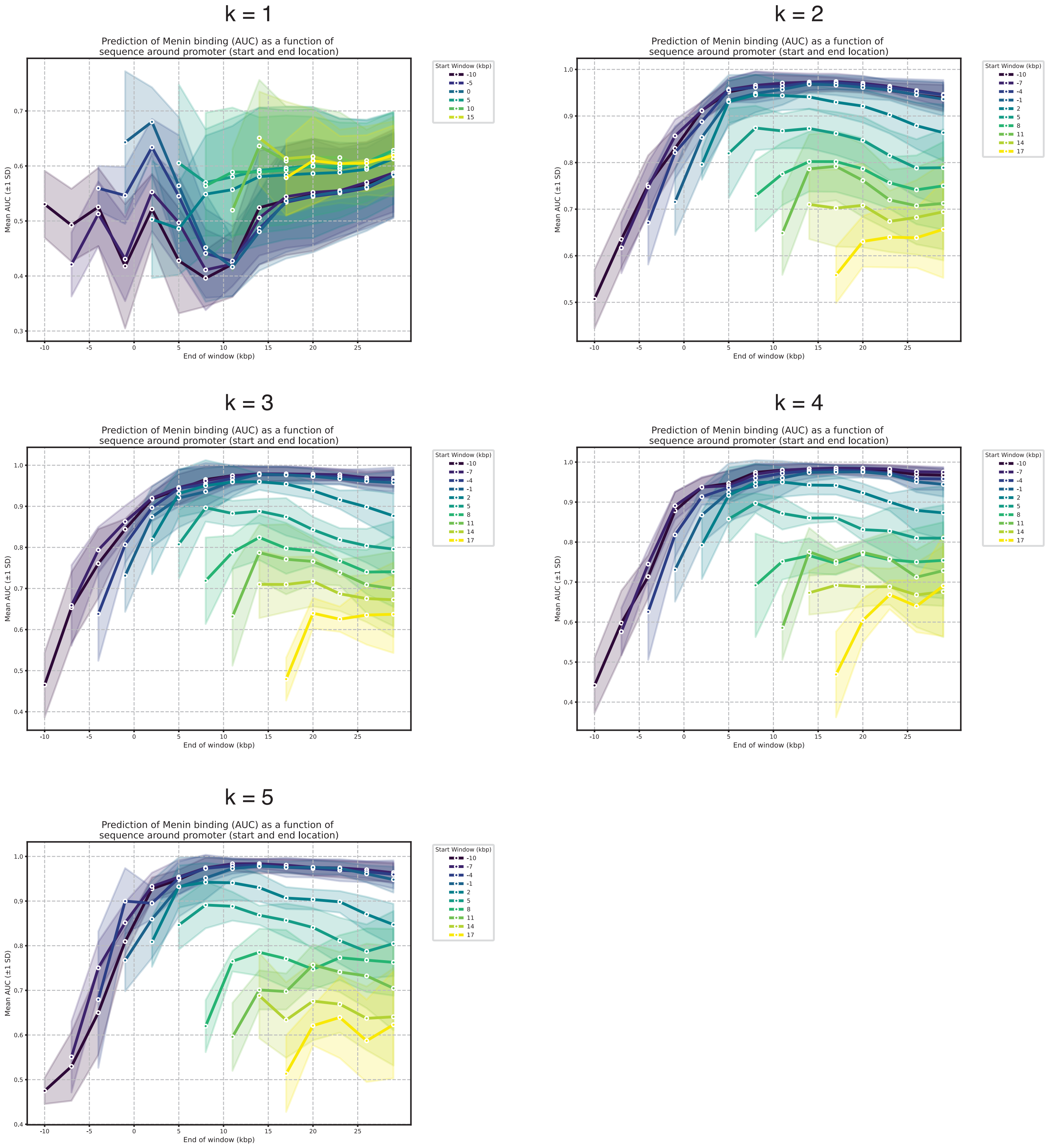


**Figure S3. Logistic regression analysis of Menin-KMT2A$^{high}$ promoters.** Logistic regression classification of Menin-KMT2A$^{high}$ and Menin$^{low}$ promoters derived from MV4-11 and OCI-AMl3 cells. Inputs consisted of DNA k-mer counts of different sizes, and each graph depicts a different k-mer size. The ROC AUC was calculated using different promoter windows (line color depicts the window start and X-axis depicts the window end). The ROC AUC was calculated using a 5-fold cross-validation on the input set; the main line represents the AUC average and the shaded area the standard deviation. ROC, Receiver operating characteristic; AUC, area under the curve.

### Supplemental Figure 4

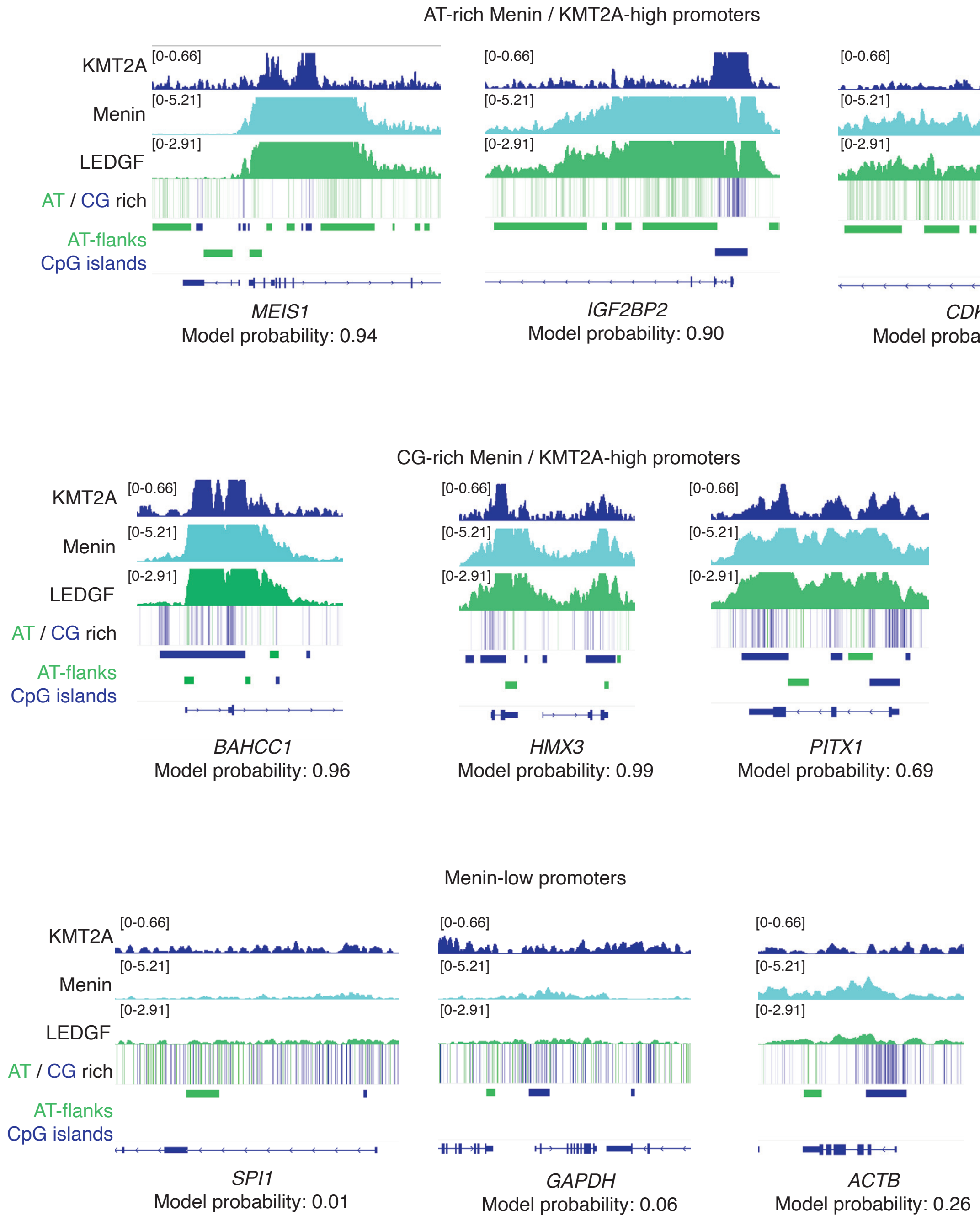


**Figure S4. Additional examples of Menin-KMT2A$^{high}$ and Menin$^{low}$ promoters.** Representative ChIP-seq tracks for KMT2A, Menin and LEDGF in MV4-11 cells at Menin-KMT2A$^{high}$ promoters with large AT-flanks (top), small AT-flanks (middle), and Menin$^{low}$ promoters (bottom).

### Supplemental Figure 5

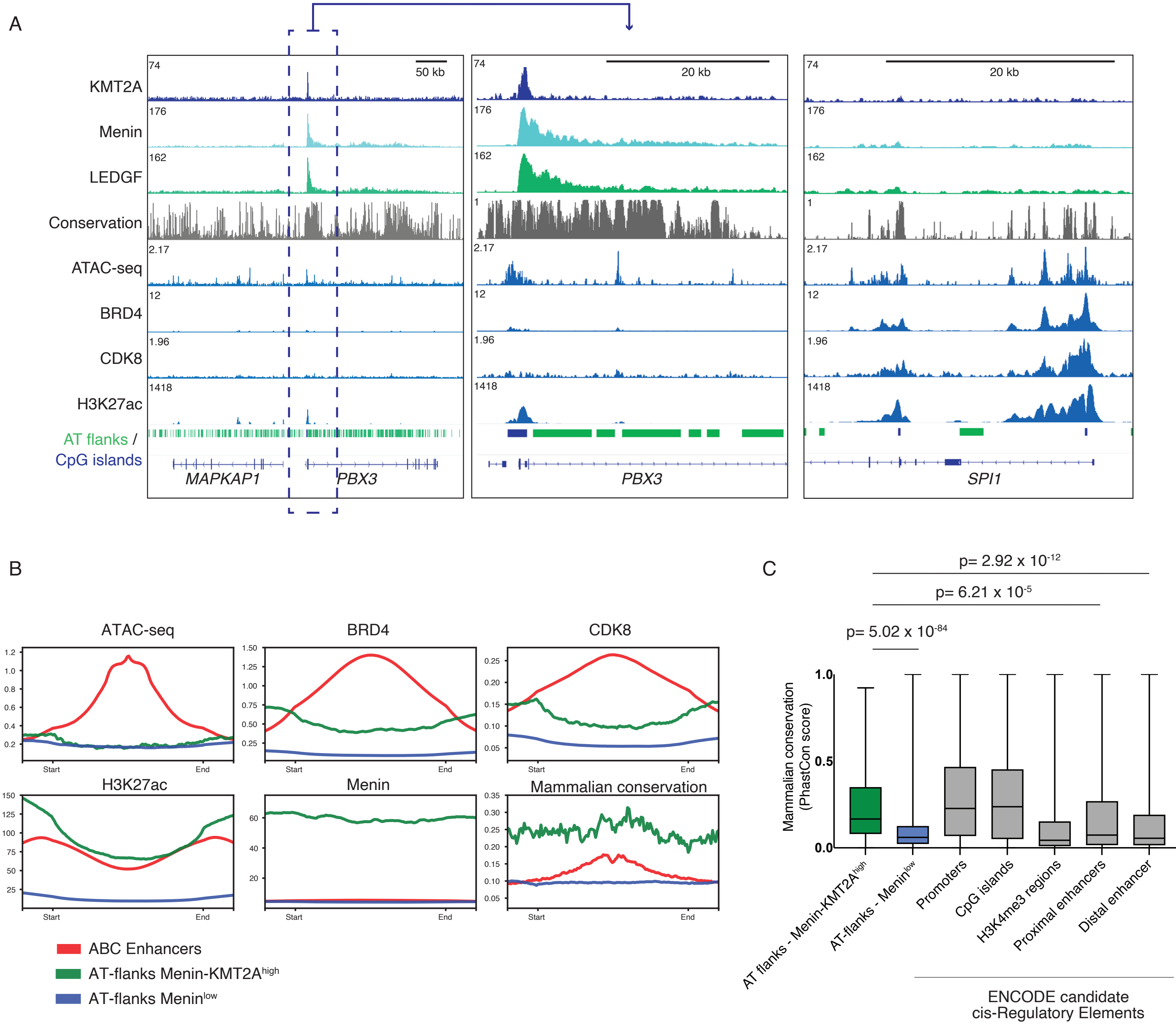


**Figure S5. AT-flanks are conserved regulatory elements distinct from canonical enhancers. a,** Mammalian conservation and ChIP/ATAC-seq tracks for KMT2A complex members (KMT2A, Menin, LEDGF) and enhancer-associated markers (ATAC-seq, BRD4, CDK8, H3K27ac) at PBX3 (Menin-KMT2A$^{high}$) and SPI1 (Menin$^{low}$). **b,** Average signal profiles of ATAC-seq, BRD4, CDK8, H3K27ac, Menin and mammalian conservation at enhancers and AT-flanks. **c,** Average mammalian conservation scores of AT-flanks compared to ENCODE candidate cis-regulatory elements. Error bars indicate s.d.; P-values were calculated using a two-tailed t-test.

### Supplemental Figure 6

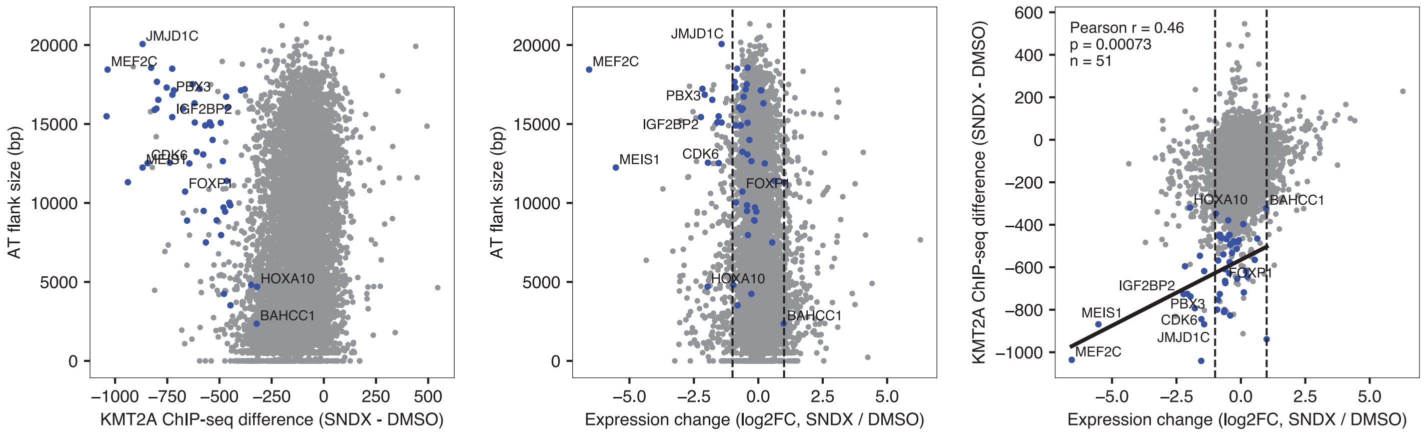


**Figure S6. Effects of Menin inhibitor treatment in KMT2A occupancy and gene expression.** Comparison between KMT2A chip-seq difference and gene expression changes of MV4-11 cells treated with Menin inhibitor for 5 days (SNDX-5613, 1 $\mu$M, 5 days). ChIP-seq coverage and AT-flank sizes were calculated using a -3 to +20 kb window. Blue dots represent Menin$^{high}$ genes defined on Figure 2. Vertical dotted lines on the middle and right panels represent log2 fold-change = 1 or -1. Data obtained from GSE196035.

### Supplemental Figure 7

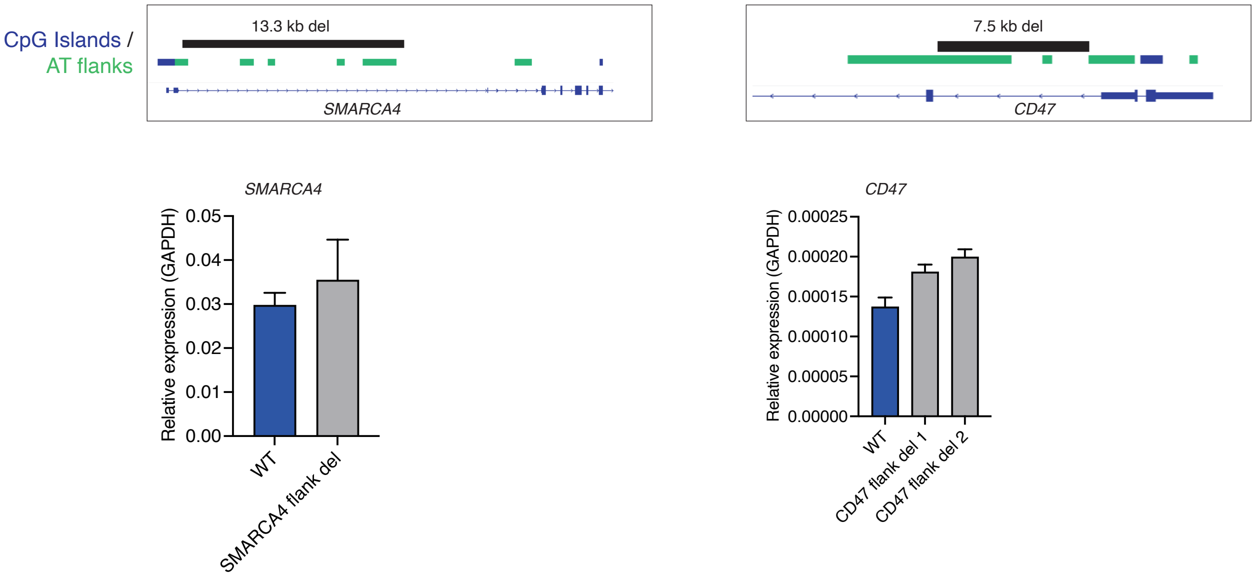


**Figure S7. Effects of flank deletion on Menin$^{low}$ promoters.** RT-qPCR analysis of *SMARCA4* and *CD47* expression following promoter flank deletion.

### Supplemental Figure 8

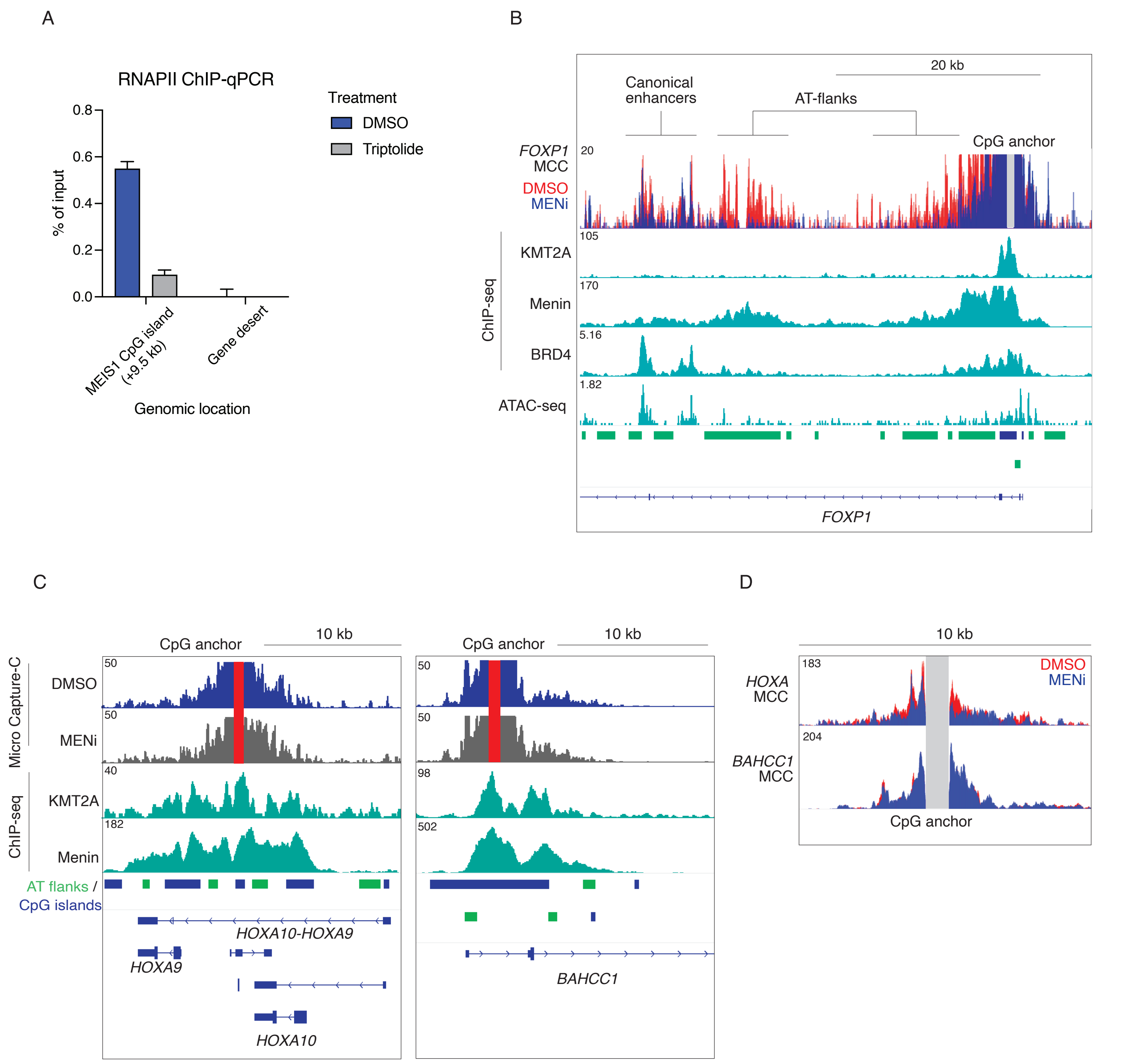


**Figure S8. Additional Micro Capture-C analysis on select Menin-KMT2A$^{high}$ targets. a,** ChIP-qPCR analysis of RNA Pol II binding to *MEIS1* CpG island (+9.5 kb) upon treatment with triptolide (500 nM, 1h). **b,** Micro Capture-C (MCC) overlay of the *FOXP1* locus in MV4-11 cells upon Menin inhibition (VTP-50469, 500 nM, 3 days). ChIP-seq tracks for KMT2A, Menin, BRD4 and ATAC-seq are shown. **c,d,** MCC analysis (c) and overlay (d) of the *HOXA* and *BAHCC1* loci upon treatment with Menin inhibitor. Capture region shown in red (c) or gray (b,d).MENi, Menin inhibitor.

### Supplemental Figure 9

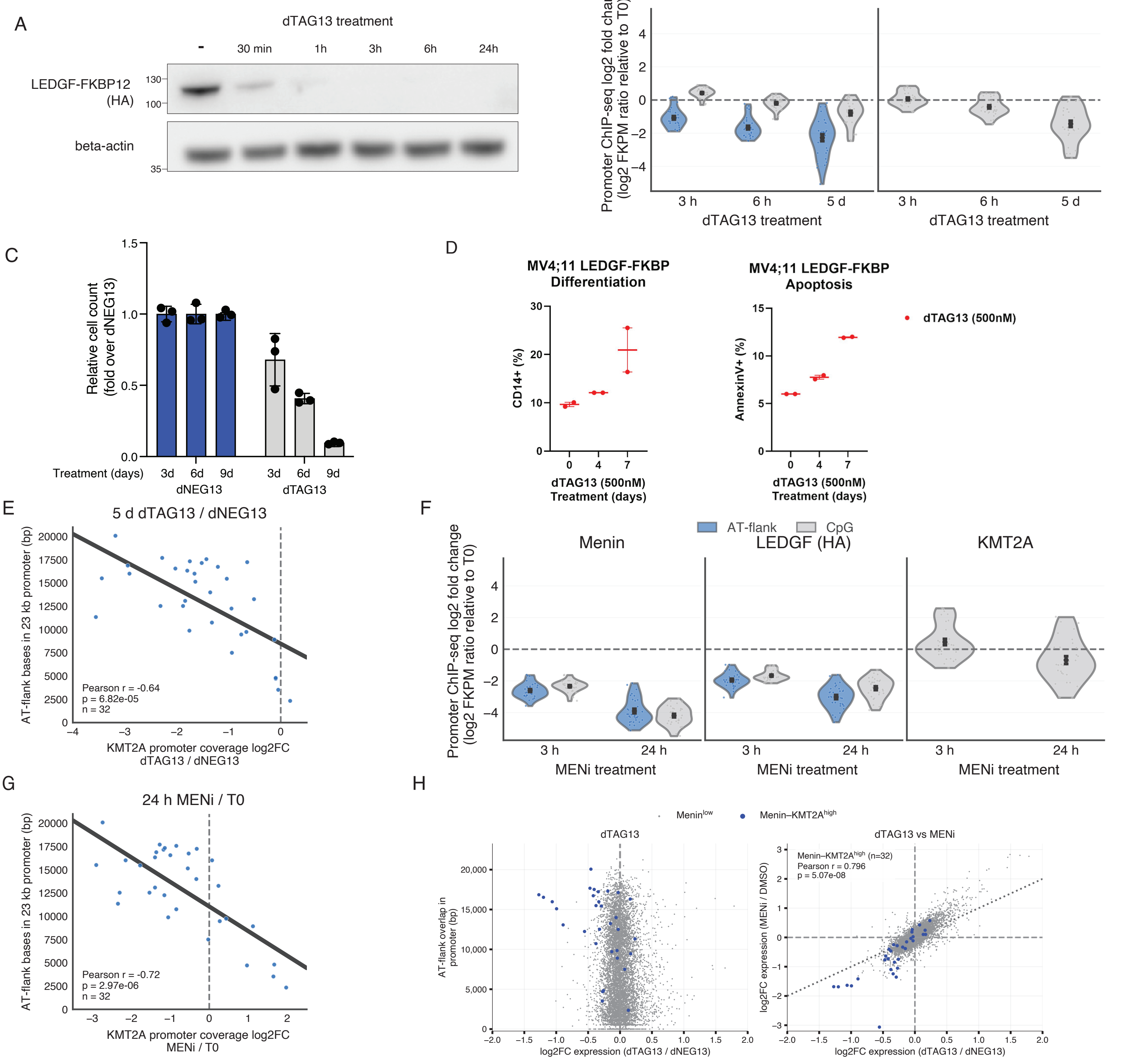


**Figure S9. Additional characterization of chromatin and transcriptional effects of LEDGF degradation and Menin inhibition in LEDGF-FKBP12-HA MV4-11 cells. a,** Time course of dTAG13-induced degradation of LEDGF measured by western blot. **b,** Log2 fold-change in KMT2A and Menin ChIP-seq coverage at CpG and AT-flanks of Menin-KMT2A$^{high}$ promoters (-3 to +20 kb) upon dTAG13 treatment. **c,** Relative cell count of cells treated with dTAG13 or dNEG13. **d,** Differentiation and apoptosis upon LEDGF degradation. **e,** Correlation between KMT2A coverage loss upon LEDGF degradation and AT-flank content at Menin-KMT2A$^{high}$ promoters. **f,** Log2 fold-change in KMT2A, Menin and LEDGF (HA) ChIP-seq coverage at CpG and AT-flanks of Menin-KMT2A$^{high}$ promoters (-3 to +20 kb) upon MENi treatment. **g,** Correlation between KMT2A coverage loss upon MENi treatment and AT-flank content at Menin-KMT2A$^{high}$ promoters. **h,** Scatter plots comparing gene-expression changes following 48-h LEDGF degradation with promoter AT-flank extent (left) and with expression changes following MENi treatment (right). MENi (Menin inhibitor): VTP-50469, 500 nM (f,g) or SNDX-5613, 50 nM (h). Error bars represent standard deviation. Specific fold-changes in KMT2A coverage at AT-flanks were not computed due to low KMT2A occpancy at these regions (b,f).

### Supplemental Figure 10

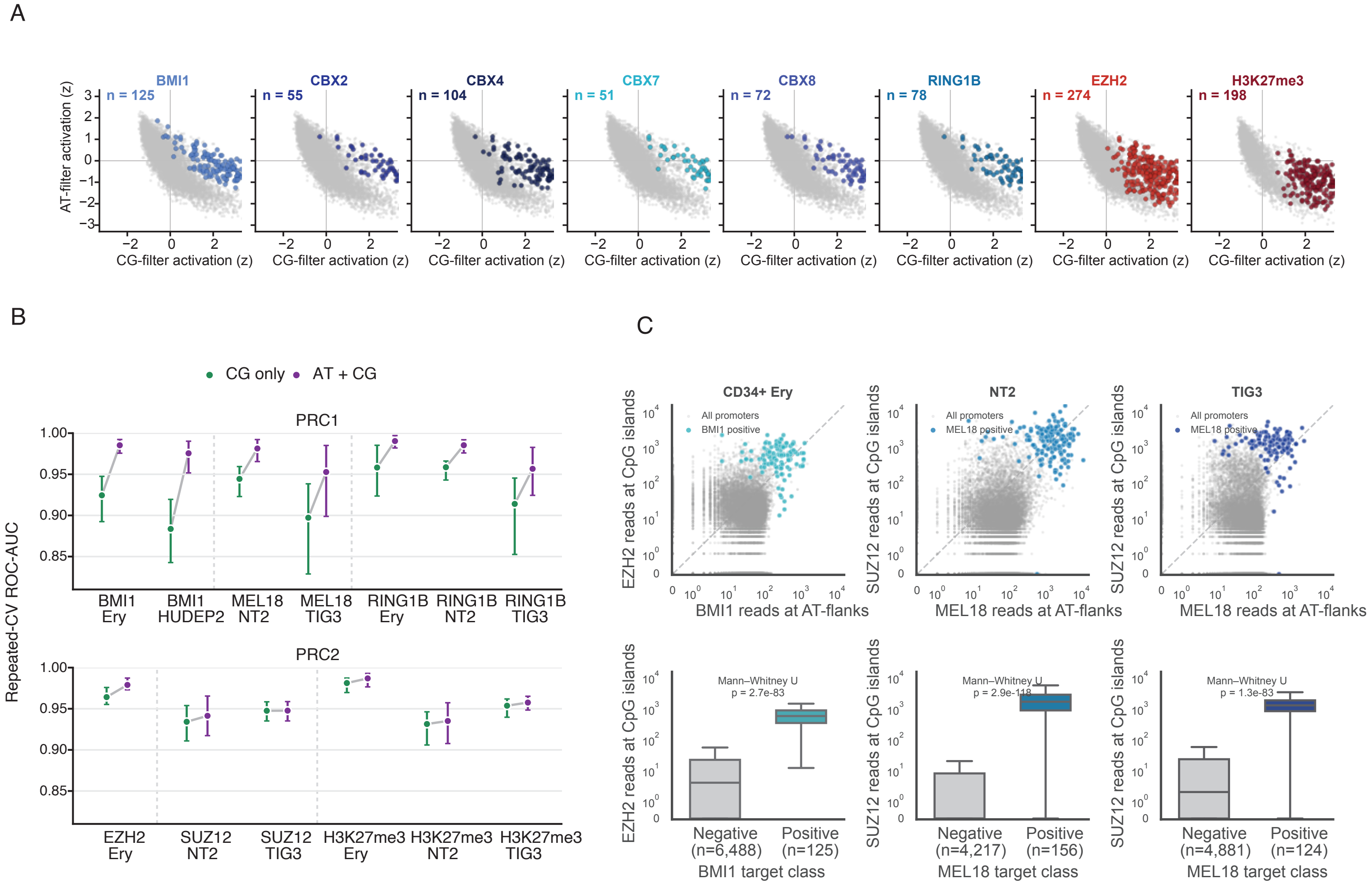


**Figure S10. Additional characterization of CpG island-AT-flank promoter architecture in Polycomb complexes. a,** Average kernel outputs for CG- and AT-rich filter across promoter regions of silenced target genes and silenced non-targets. Target promoters are shown in shades of blue for cPRC1 components and shades of red for PRC2 components and H3K27me3. Non-target promoters are shown in gray.. **b,** Logistic regression performance using CG-filter activation alone or combined CG- and AT-filter activations in CD34+-derived erythrocytes (Ery), HUDEP2, NT2, and TIG3 cells. Points indicate mean ROC-AUC from repeated stratified cross-validation; error bars indicate the 2.5th–97.5th percentiles. **c,** Comparison of CUT&RUN and ChIP-seq coverage for cPRC1 components at AT-flanks and PRC2 components at CpG islands across different publicly available datasets (top row), and PRC2 coverage at silenced BMI1/MEL18$^{high}$ (positive) and BMI1/MEL18$^{low}$ (negative) promoters (bottom row). Boxes indicate the interquartile range with median; whiskers extend to 1.5× IQR.

### Supplemental Figure 11

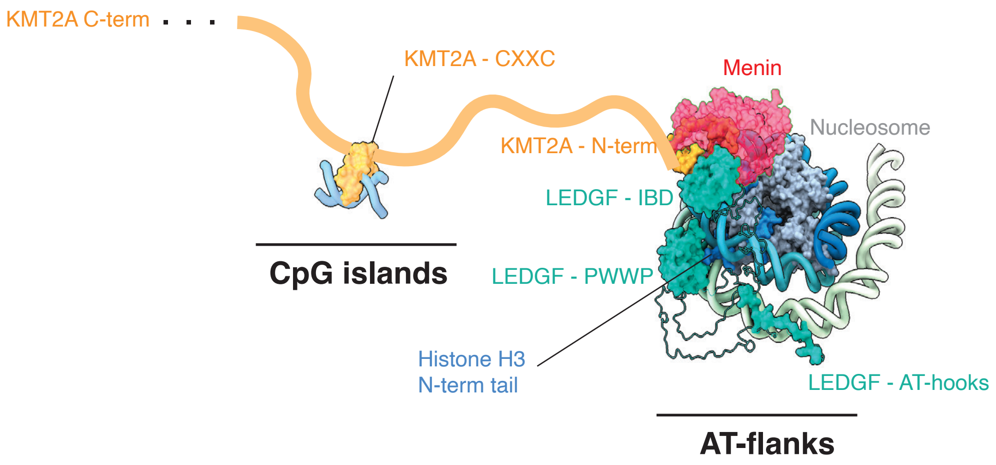


**Figure S11. Schematic model of KMT2A, Menin and LEDGF engagement with AT-flanks and CpG islands.** Structurally inspired composite model assembled from experimental structures of the KMT2A–Menin–LEDGF IBD complex (X-ray crystallography, PDB 3U88), the LEDGF PWWP–nucleosome complex (cryo-EM, PDB 6S01) and the Menin–nucleosome complex (cryo-EM, PDB 8GPN), together with AlphaFold3 predictions of the LEDGF AT-hook bound to A/T-rich DNA and of the KMT2A CXXC domain bound to CpG-containing DNA. LEDGF domain linkers not resolved in the source structures were built with MODELLER. The AT-hook position relative to the nucleosome is inferred rather than experimentally constrained, as the AT-hook was not resolved in PDB 6S01 and that structure used a Widom 601 construct whose flanking DNA is not A/T-rich. The relative orientation of the LEDGF PWWP and IBD domains has not been experimentally determined.
